# Ligand-mediated structural dynamics of a mammalian pancreatic K_ATP_ channel

**DOI:** 10.1101/2022.03.02.482692

**Authors:** Min Woo Sung, Camden M. Driggers, Barmak Mostofian, John D. Russo, Bruce L. Patton, Daniel M. Zuckerman, Show-Ling Shyng

## Abstract

Regulation of pancreatic K_ATP_ channels involves orchestrated interactions of channel subunits, Kir6.2 and SUR1, and their ligands. How ligand interactions affect channel conformations and activity is not well understood. To elucidate the structural correlates pertinent to ligand interactions and channel gating, we compared cryo-EM structures of channels in the presence and absence of pharmacological inhibitors and ATP, focusing on channel conformational dynamics. We found pharmacological inhibitors and ATP enrich a channel conformation in which the Kir6.2 cytoplasmic domain is closely associated with the transmembrane domain relative to one where the Kir6.2 cytoplasmic domain is extended away into the cytoplasm. This conformation change remodels a network of intra and inter-subunit interactions as well as both the ATP and PIP_2_ binding pockets. The structures resolved key contacts between the distal N-terminus of Kir6.2 and SUR1’s ABC module involving residues implicated in channel function. A SUR1 residue, K134, is identified to directly contribute to the PIP_2_ binding pocket. Molecular dynamics simulations revealed two Kir6.2 residues, K39 and R54, that mediate both ATP and PIP_2_ binding, suggesting a mechanism for competitive gating by ATP and PIP_2_.

## Introduction

Pancreatic ATP-sensitive potassium (K_ATP_) channels functionally couple glucose metabolism to insulin release and are crucial for glucose homeostasis (Ashcroft, 2005; Nichols, 2006). Structurally, the pancreatic K_ATP_ channel is an octameric complex composed of two distinct integral membrane proteins (Clement et al., 1997; Inagaki et al., 1997; Lee et al., 2017; Li et al., 2017; Martin et al., 2017b; Shyng and Nichols, 1997). A tetrameric core of Kir6.2 subunits form the central transmembrane pore of the channel. A coronal array of four sulfonylurea receptor 1 (SUR1) subunits surrounds the channel core, each SUR1 companioned with one Kir6.2 subunit. Genetic mutations of these subunits that dysregulate K_ATP_ channel activity are causes of neonatal diabetes (gain of function) and congenital hyperinsulinism (loss of function) (Ashcroft, 2005). K_ATP_ channels harbor multiple distinct and antagonistic binding sites for their primary physiological regulators, intracellular ATP and ADP, which close the ion channel through a binding site in Kir6.2, but open the channel through Mg-dependent binding sites on SUR1 (Nichols et al., 1996; Puljung, 2018). In addition, channel activity is operationally governed by binding sites for specific membrane phospholipids, most particularly PIP_2,_ which directly promote opening as well as antagonize the ATP inhibition at the Kir6.2 binding sites (Nichols et al., 1996). Finally, the pancreatic K_ATP_ channel is the drug binding target for sulfonylurea and glinide anti-diabetic medications, which inhibit channel activity and thus stimulate insulin secretion (Gribble and Reimann, 2003). The long held principal objective of K_ATP_ channel research has been understanding the protein dynamics by which these several ligand interactions, separately and in concert, ultimately determine levels of K_ATP_ channel activity, and hence control insulin release.

CryoEM structures of K_ATP_ channels have provided direct insights into the structural mechanisms of ligand recognition and gating regulation. Previously, we reported comparative cryoEM structures for pancreatic K_ATP_ channels in the absence of ligands (apo); in the presence of ATP; and in the combined presence of ATP with alternative pharmacological inhibitors, glibenclamide, repaglinide, or carbamazepine (Martin et al., 2019). The study found all pharmacological inhibitors occupy a common binding pocket located within SUR1. Further, we found SUR1’s common drug binding site lies adjacent to the deep binding site for the distal end of the extended Kir6.2 N-terminal peptide, which courses through the prominent cleft between the two halves of the ABC (ATP Binding Cassette) module of SUR1. The findings offered mechanistic insight into how distinct pharmacological inhibitors inhibit channel activity and also facilitate channel assembly by stabilizing the interaction between Kir6.2 N-terminus and SUR1. However, it was noted during image analysis that each dataset in the study possessed considerable conformational heterogeneity, suggesting comparative analyses within datasets may further illuminate channel structural dynamics relevant to ligand binding and gating.

Here, we show results from reprocessing of cyroEM datasets previously reported, focusing on conformational analysis and augmented by molecular dynamics (MD) simulations. Most notably, we found that the cytoplasmic domain (CTD) of Kir6.2 adopted two distinct conformations. In one, the CTD is tethered close to the membrane (Kir6.2-CTD-up). In the other, the CTD is counterclockwise corkscrewed away from the membrane, towards the cytoplasm (Kir6.2-CTD-down). Across structure datasets, the ratio of CTD-up versus CTD-down conformations strongly correlated with the occupation of inhibitory ligand binding sites. In addition, drug binding and CTD conformation were associated with significant structural reorganization at the ATP and PIP_2_ binding sites, and at domain interfaces within and between subunits, suggesting ligands act as molecular glues to stabilize the channel in the Kir6.2-CTD-up conformation. Of further importance, improved cryoEM maps and functional analysis revealed that binding of the activating ligand PIP_2_ leading to channel gating involves an interaction with SUR1 via a positively charged lysine residue in TMD0. MD simulations uncovered further residues that participate in both ATP and PIP_2_ binding. The findings implicate mechanisms by which SUR1 enhances Kir6.2 PIP_2_ sensitivity, and explain the manner by which ATP and PIP_2_ compete to control K_ATP_ channel gating. Finally, they provide a consistent structural framework for understanding how mutations, now observed to affect key protein-protein and protein-ligand interfaces, disrupt channel function and cause disease.

## Results

### K_ATP_ channel conformation analysis

Focused 3D classification on the Kir6.2 tetramer core plus one SUR1 subunit (denoted K_4_S hereinafter) following symmetry expansion and signal subtraction (Scheres, 2016) was performed on our previously published five datasets: apo, ATP only, carbamazepine and ATP (CBZ/ATP), glibenclamide and ATP (GBC/ATP), repaglinide and ATP (RPG/ATP) (Martin et al., 2019) (see Methods). This strategy was employed to circumvent alignment difficulty due to flexible SUR1 (Fig.S1). The analysis revealed two major K_4_S conformations: Kir6.2-CTD-up and Kir6.2-CTD-down, wherein the Kir6.2-CTD was alternatively located closer to, or further from, the Kir6.2 membrane spanning channel domains, respectively. More particularly, translation of the CTD between up and down conformations further involved a rotation, together comprising a corkscrew movement wherein the CTD (from an extracellular point of view) was rotated clockwise in Kir6,2-CTD-up conformation relative to Kir6.2-CTD-down (Fig.1). The CTD-up and CTD-down conformations appear qualitatively simialr to the T(tense)-state and R(relaxed)-state previously reported by others using a fusion SUR1-Kir6.2 protein (Wu et al., 2018). Within both the CTD-up and CTD-down conformations, rocking and rotation of the Kir6.2-CTD was discernable using Relion 3 multibody refinement principal component analysis (Nakane et al., 2018). The heterogeneity was greater in the CTD-down than the CTD-up population of particles (Fig.S2), consistent with an increase in CTD mobility when detached from the membrane. Similar Kir6.2-CTD dynamics were observed using cryoSPARC 3D variability analysis (Punjani and Fleet, 2021).

**Figure 1.**
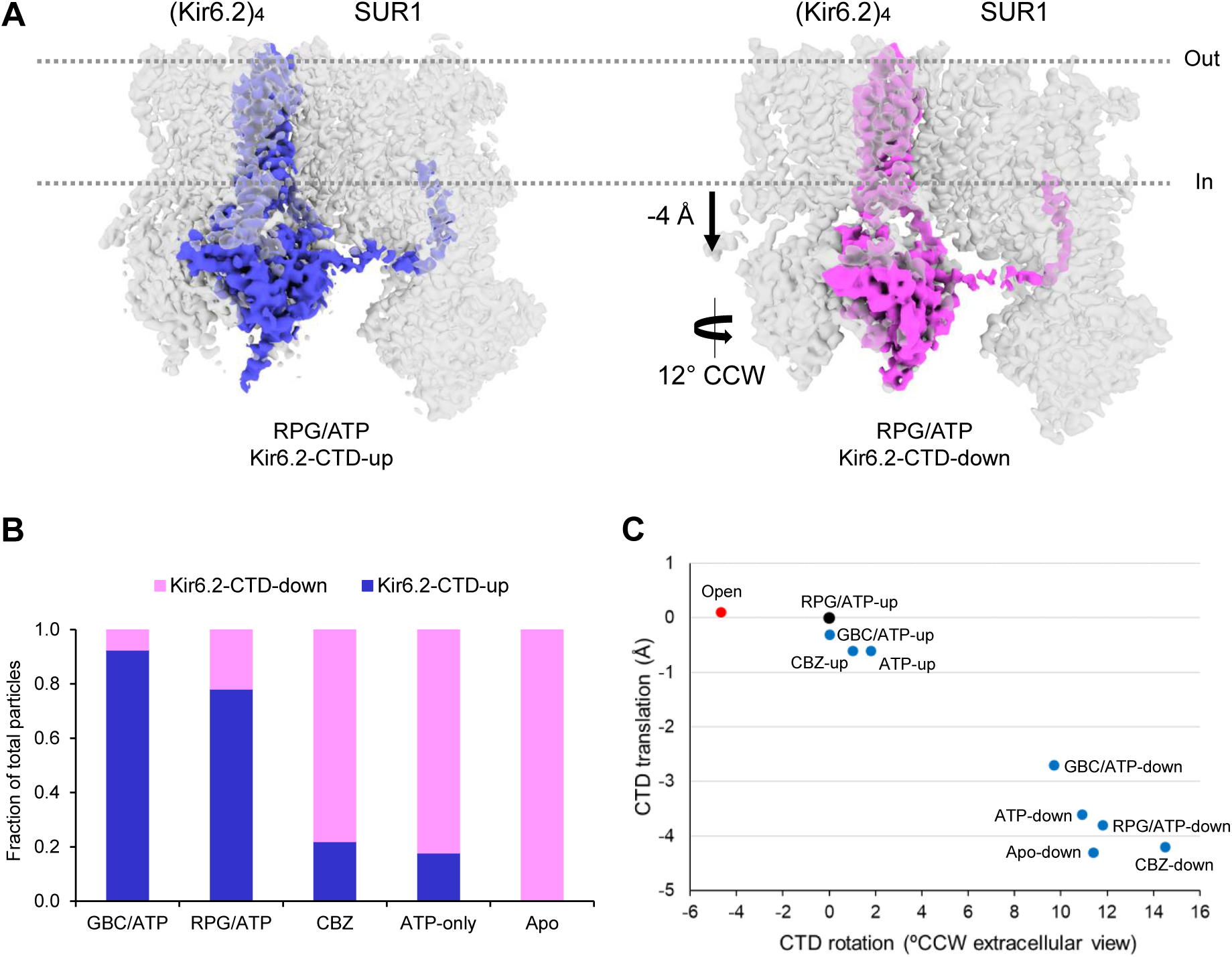
Two distinct conformations of the Kir6.2-CTD in RPG/ATP bound K_ATP_ channels. **(A)** In the Kir6.2-CTD-down conformation, the CTD is translocated away from the inner lipid bilayer towards the cytoplasm by ∼4 Å and rotated counterclockwise by 12º (viewed from the extracellular side) compared to Kir6.2-CTD-up conformation. The distance between the G-loop gate (center of mass of G295-Cα of all four Kir6.2 subunits) and the helix bundle crossing gate (center of mass of F168-Cα of all four Kir6.2 subunits). The CTD’s rotational angles were estimated by aligning the structure onto the RPG CTD-up reference model TM domain (residues 55-175), and calculating a dihedral angle between K338-Cα of Chain A of Kir6.2 and the center of mass of G245-Cα. **(B)** Fraction of particles in Kir6.2-CTD-up and Kir6.2-CTD-down in K_ATP_ channels bound to different ligands or in apo state. **(C)** Variations in the extent of CTD translation and rotation observed in all datasets (blue circles) using RPG/ATP-Kir6.2-CTD-up as reference structure (black circle). Translation and rotation of the CTD from a human open K_ATP_ structure (Zhao and MacKinnon, PDB: 7S5T) relative to the RPG/ATP Kir6.2-CTD-up reference structure is included for comparison (red circle). CTD translation away from the membrane is shown in negative value.

Both the Kir6.2-CTD-up and Kir6.2-CTD-down conformations are observed in the GBC/ATP, RPG/ATP, CBZ/ATP and ATP-only datasets; however, relative abundance of the two conformations varies in the different liganded states (Fig. 1B, Table S1). The GBC/ATP and RPG/ATP datasets have the highest percentages of particles in the CTD-up conformation, with 92.5% and 71.2%, respectively. The CBZ/ATP dataset, which only has CBZ density in SUR1 but no ATP density in Kir6.2 likely due to lower concentrations of ATP used during sample preparation (see Methods) has a significantly lower percentage (22%) of particles in the CTD-up conformation, which is comparable to the ATP only dataset of 17.8%. In the absence of added ligands, or the apo state, no Kir6.2-CTD-up, only Kir6.2-CTD-down conformation was observed. These findings show that Kir6.2-CTD exists largely in two discrete conformations and the distribution of the two states is correlated with binding of inhibitory ligands, implying a mechanistic role for the conformations. The RPG/ATP dataset gave the highest resolution maps for both the Kir6.2-CTD-up and CTD-down conformations (3.4 Å and 3.6 Å, respectively; Fig.S1). These maps have improved quality compared to our previously published structure (Martin et al., 2019), allowing us to reevaluate ligand and protein densities that were previously ambiguous. We therefore focused on this dataset for structural analyses hereinafter.

In the RPG/ATP state, the predominant SUR1 conformation is arranged like a propeller when symmetrized as described in our previous study (Martin et al., 2019). In addition, a minor conformation (∼27%; Fig.S1) showing a large clockwise rotation of SUR1 towards the Kir6.2 tetramer (viewed from the extracellular side) was identified. This conformation is qualitatively similar to the quatrefoil conformation previously reported in the MgATP/MgADP-bound, NBDs-dimerized Kir6.2-SUR1 fusion channel structure (Lee et al., 2017), and our recently reported quatrefoil-like Kir6.1-SUR2B vascular K_ATP_ channel structure bound to GBC and ATP with separate NBDs (Sung et al., 2021). The overall map resolution of this minor class is ∼7 Å (Fig.S1A), which precludes detailed structural analysis. Nonetheless, it implies that even in the presence of RPG and ATP, a large rotation of SUR1 resembling that seen in NBDs dimerized SUR1 quatrefoil conformation occurs, albeit much less frequently. Heterogeneity of SUR1 within the dominant propeller conformation with more subtle rotations of SUR1’s ABC core around the Kir6.2 tetramer is also observed whether the Kir6.2-CTD is up or down. We focused on refining SUR1 structures in the dominant Kir6.2-CTD-up class. Particles with eigenvalues between 5 and 20 or -5 and -20 along eigenvector 1 were refined separately to generate two maps we referred to as SUR1-in and SUR1-out conformations at 3.9 Å and 3.8 Å (Fig.S3), respectively, for model building (Table S2) and structural analysis.

### Comparison of different K_ATP_ conformations

The changing conformations of Kir6.2-CTD and SUR1 remodeled subunit and domain interfaces as well as protein-ligand interactions. Comparing CTD-down to the CTD-up conformation, the Kir6.2-CTD is translated down into the cytoplasm by ∼4 Å along an axis perpendicular to the embedding membrane, and simultaneously counterclockwise (CCW) rotated by 12º about that axis viewed from the extracellular side (Fig.1). The altered location of Kir6.2-CTD remodeled the interfacial (IF) helix (also called the slide helix, herein taken to include G53-D65) in the Kir6.2 N-terminus, and also the C-linker (herein taken to include H175-L181) by which inner helix M2 interacts with the CTD. Specifically, the IF helix as composed in the CTD-down conformation formed a continuous helix that extended towards the neighboring Kir6.2 subunit. In contrast in the CTD-up conformation, the IF helix adopted a 3_10_ helix characteristic (Vieira-Pires and Morais-Cabral, 2010) wherein a directional kink at D58 demarcated the helix into N-terminal and C-terminal halves, with the N-terminal half pivoted towards SUR1 instead of along the adjacent Kir6.2 subunit. Respecting the C-linker, while in the CTD-down conformation, the C-linker was fully unraveled into a loop, in which a key PIP_2_ interacting residue R176 (Baukrowitz et al., 1998; Shyng and Nichols, 1998) was distant from the membrane and incapable of direct PIP_2_ interaction, in the CTD-up conformation, the C-linker formed a helical structure that participated in membrane PIP_2_ binding. In comparisons between the SUR1 propeller in and out structures (Fig.S1, S3), the ABC module in the SUR1-in structure is rotated clockwise closer to a neighboring Kir6.2 (viewed from the extracellular side). In this rotated position, the SUR1-L0 loop was pulled away from its interaction with SUR1’s direct Kir6.2 subunit partner. Specific molecular changes at the subunit and domain interfaces and ligand binding sites in different conformations are described below.

### Intra-Kir6.2 and inter Kir6.2-Kir6.2 interactions

Close inspection of the Kir6.2 CTD-up and CTD-down structures revealed two salt bridges in the CTD-up conformation, formed between D58 and R206 from adjacent Kir6.2 subunits and between R177 and D204 in the same subunit, which became fully disrupted in the CTD-down conformation (Fig.2). In combination, the two ion pairs bound together three structural elements that occupy the transitional space between the membrane spanning and cytoplasmic domains of Kir6.2. Residues D204 and R206 are both at the start of a loop at the top of the CTD that connects βD and βE (DE loop) (Martin et al., 2017b). In the CTD-up conformation, D204:R177 linked the DE loop to the C-linker, which connects to Kir6.2 TM helix 2; and R206:D58 linked the DE loop to the IF helix of the adjacent Kir6.2 subunit. Both salt bridges were eliminated in the CTD-down structures. Separation of D204 from R177 in CTD-down was accompanied by uncoiling and extension at the end of the C-linker helix including R177. In contrast, separation of R206 from D58 was accompanied by a significant straightening of the kink in the IF helix and reorientation of the continuing chain, which connects to the base of the KNtp (Fig.2).

**Figure 2.**
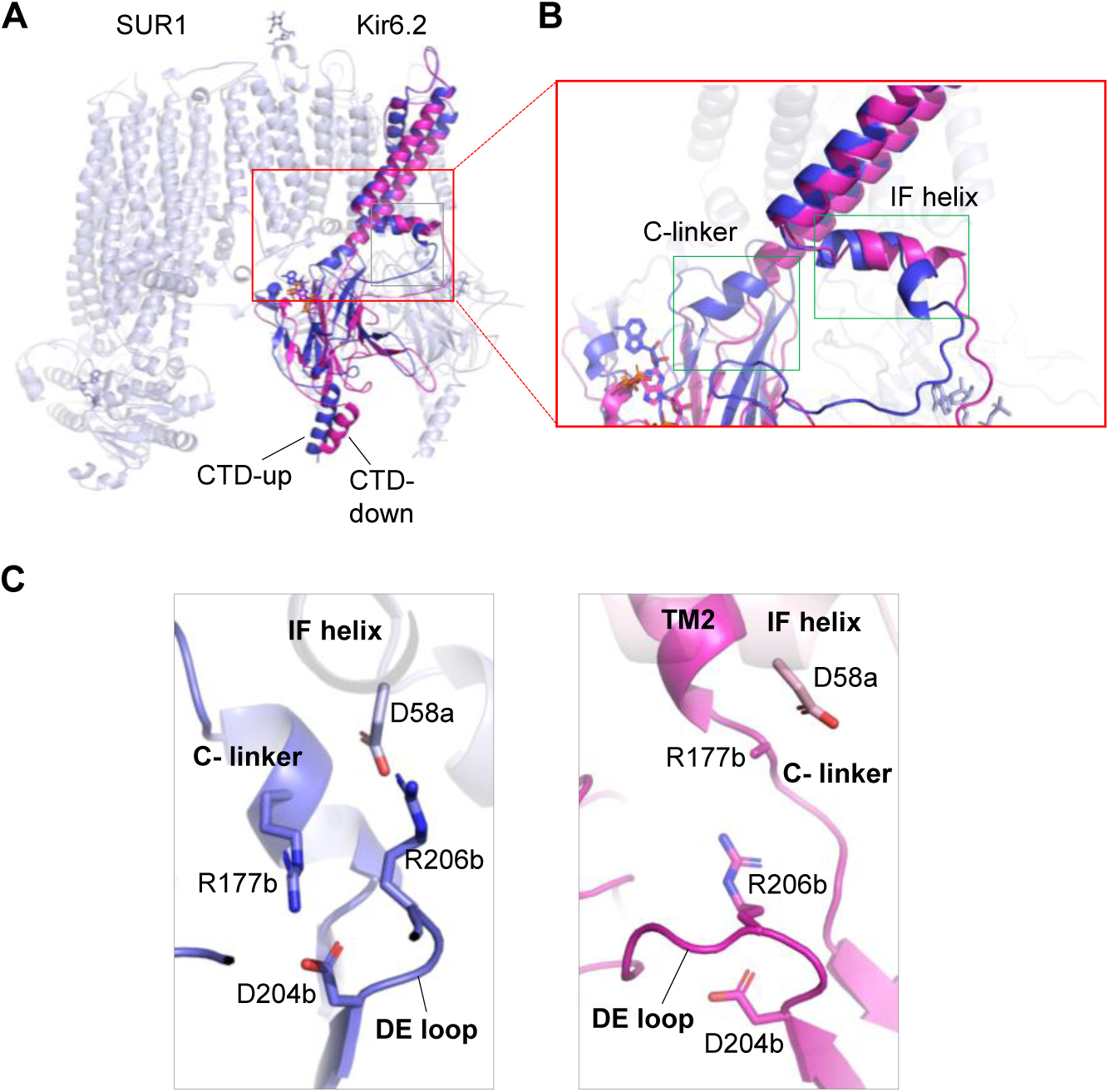
Structural comparison between RPG/ATP CTD-up and CTD-down conformations. **(A)** Superposition of the Kir6.2-CTD-up (blue) and CTD-down (magenta) structures. The red boxed region shows significant secondary structural difference at the slide helix and the C-linker. **(B)** Close-up view of the structural difference at the slide helix and the C-linker in the two conformations. **(C)** Close-up view of the molecular interactions between D58 and R206 of neighboring Kir6.2 subunits and R177 and D204 from the same Kir6.2 subunit in the Kir6.2-CTD-up conformation (left), and loss of these ion pair interactions in the Kir6.2-CTD-down conformation (right).

Consistent with the notion that these labile salt bridges have critical roles in channel activation, previously published functional studies have implicated the participating Kir6.2 residues in channel regulation. D58 was previously suggested to be involved in anchoring Kir6.2 CTD to the TM domain through results of targeted mutagenesis (Li et al., 2013). Our findings resolve the salt bridge partnership with R206 in the neighboring Kir6.2 polypeptide, and further reveal such tethering is labile and dynamically incorporated into the conformational changes of ion channel activation. Consolidating this view, a critical role for R206 has been separately implicated in the gating interactions with PIP_2_ that govern channel opening, through scanning mutagenesis investigations of positive residues involved in effecting bound PIP_2_, wherein mutation R206A was found to abolish PIP_2_ response and thus diminish channel activity (Li et al., 2013; Lin et al., 2005; Shyng et al., 2000). Similar to R206A, mutation R177A also abolishes or greatly attenuates channel activity by dimishing PIP_2_ response (Li et al., 2013; Shyng et al., 2000). Thus, in corroborating earlier functional studies, our structural findings here elucidate key molecular interactions that hold the Kir6.2-CTD close to the membrane in position to interact with membrane-bound PIP_2_ for channel opening. These insights are directly relevant to human health. Mutations of each of the four residues in the above interactions have been identified in congenital hyperinsulinism, a disease caused by loss of function of K_ATP_ channels. These include D58V (De Franco et al., 2020), R177W (Arya et al., 2014), D204E (Pinney et al., 2008), and R206H (Boodhansingh et al., 2019). Our structures here provide a mechanistic illustration of how perturbation of residues involved in conformational dynamics cause loss of channel function and hyperinsulinism.

### Kir6.2 and SUR1 interactions

Our previous study of pancreatic K_ATP_ channel structure suggested that a key regulatory interface through which SUR1 controls Kir6.2 channel activity is formed by the extended N-terminus of Kir6.2 (residues 1-30; referred to as KNtp) that we identified deeply inserted within its SUR1 subunit partner, wherein the KNtp is located within the SUR1 ABC transporter module (Martin et al., 2019). We showed in particular that the KNtp is located between the two transmembrane helix bundles (TMBs) of SUR1 ABC module, and adjacent to the drug binding pocket of GBC, RPG, and CBZ. More detailed structural analysis was hindered by the resolution for the density of KNtp in those published cryoEM maps. The additional analysis methods applied in our current study yielded clear, contiguous densities and produced significantly improved maps including specific interactions between residues in KNtp and SUR1 (Fig.3). In the distal part of KNtp, which lies deep in the SUR1 ABC core cavity, Kir6.2-R4 is in bonding position with SUR1-T1139 and N1301. In the middle section of KNtp, which is near the entrance to the SUR1 ABC core cavity, Kir6.2-L17 interacts with R826 of NBD1 and G1119 and N1123 of TMB2. In the proximal end of KNtp, cryoEM density corresponding to residues P24, Y26 and R27 comes into close contact with cryoEM density correponding to SUR1’s NBD1-TMD2 linker around residue S988. In the Kir6.2-CTD down structure, interactions at the distal and middle segments of KNtp with SUR1 remain largely unchanged; however, the proximal section of KNtp is significantly further away from the NBD1-TMD2 of SUR1 (Fig.3D). In a recent Kir6.1-SUR2B cryoEM structure we showed that the NBD1-TMD2 linker has a role in regulating MgADP-dependent interactions between SUR2B-NBD2 and Kir6.1-CTD (Sung et al., 2021). Our structures presented here reveal an additional contact between NBD1-TMD2 linker and KNtp. Whether this contact and changes at this interface in different conformations have functional significance warrants future investigation. As reported in our previous publication (Martin et al., 2019), the cryoEM density of KNtp is the strongest in the GBC/ATP, RPG/ATP, and CBZ/ATP datasets, followed by ATP only, and is the weakest in the apo state, indicating inhibitory ligands stabilize the KNtp-SUR1 interface. Deletion of KNtp is known to increase channel open probability (Babenko et al., 1999; Koster et al., 1999; Reimann et al., 1999), while immobilizing KNtp in the SUR1 ABC core cavity via engineered crosslinking between Kir6.2-L2C and SUR1-C1142 inhibits channel activity (Martin et al., 2019). Worth noting, mutations L2P (Alkorta-Aranburu et al., 2014), R4C/H, L17P, R24C, R27C/H in Kir6.2 (De Franco et al., 2020) as well as R826W (de Wet et al., 2008) and N1123D (Suzuki et al., 2007) of SUR1 have been reported in neonatal diabetes or congenital hyperinsulinism, which further underscores the importance of this interface in channel gating

**Figure 3.**
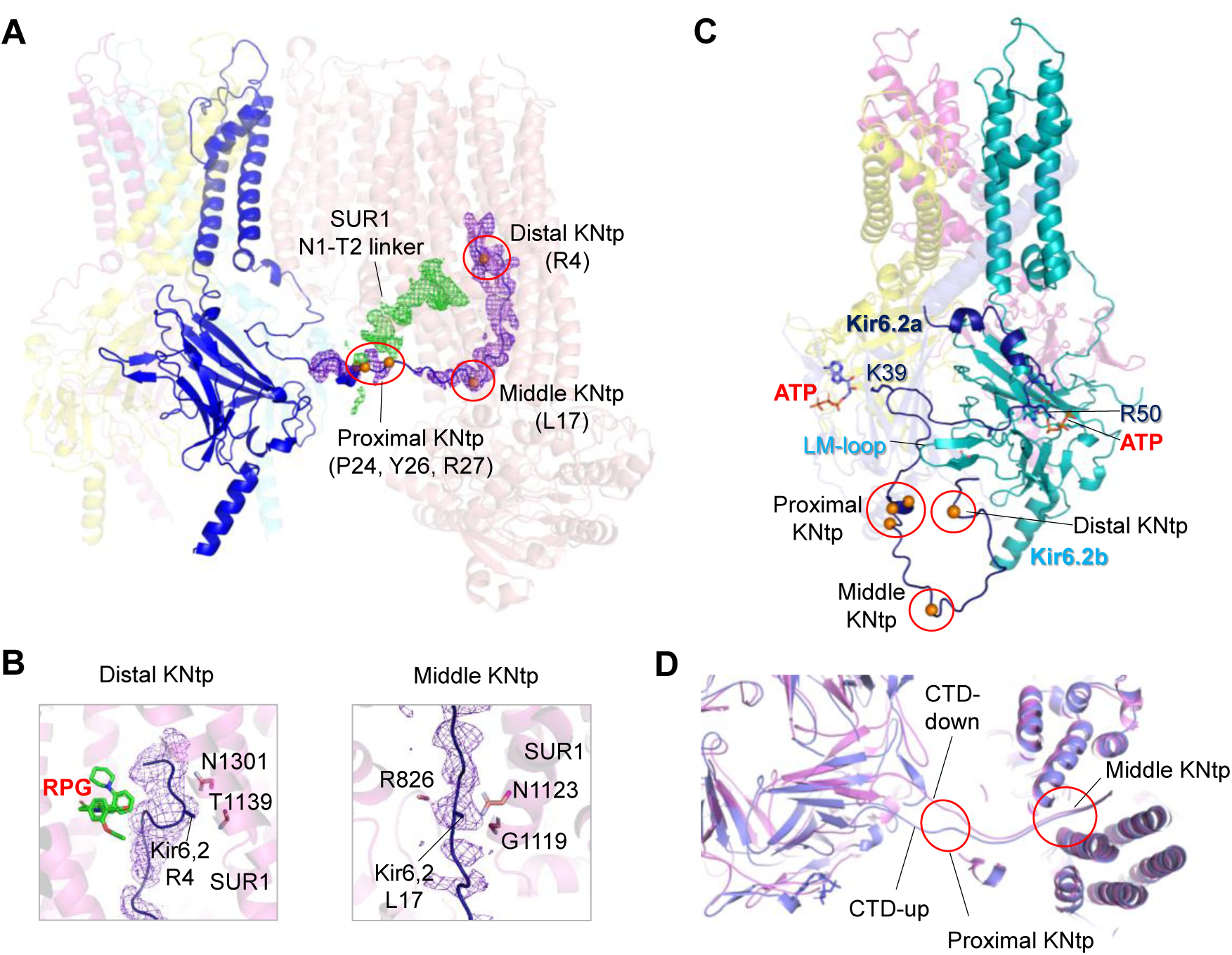
KNtp and SUR1 interface. **(A)** KNtp (Kir6.2 aa 1-30) interacts with SUR1 at multiple sites. The distal portion of KNtp located deep in the cavity of the SUR1-ABC core module. The middle section of KNtp lies near the entrance of the cavity. The proximal part (i.e. C-terminal part) of KNtp near P24, Y26 and R27 comes into contact with SUR1 N1-T2 linker density near S988. Residues in KNtp are shown as orange spheres. **(B)** *Left*: Close-up view of distal KNtp showing interaction of R4 with SUR1 T1139 and N1301 in TMD2 and proximity to bound RPG. *Right*: Close-up view of middle KNtp showing interaction of L17 with SUR1 R826, G1119 and N1123. **(C)** KNtp viewed without SUR1, showing its interconnectivity with two ATP binding sites and the LM-loop in the CTD of a neighboring Kir6.2. **(D)** Superposition of Kir6.2-CTD-up and CTD-down structures viewed from the extracellular side showing divergence of proximal KNtp.

In addition to KNtp forming interactions with the SUR1-ABC module, regions C-terminal to KNtp in the Kir6.2-N terminal domain is also intimately involved in protein-protein and protein-ligand interactions. First, in the CTD-up structure, Kir6.2 R31-R34 is close to the short loop that connects βL and βM (LM loop) (Martin et al., 2017b) of the neighboring subunit (Fig.3C). A mutation D323K in the LM loop has been shown to disrupt ATP inhibition (Brennan et al., 2020). Second, further downstream K39 has its sidechain oriented towards ATP bound to the neighboring Kir6.2 on the other side. In this way, the Kir6.2 N-terminus is connected simultaneously to two ATP binding pockets. Dynamic movement of the KNtp between the CTD-up and CTD-down conformations will likely impact the interactions of downstream Kir6.2-N terminal domain with neighboring subunits on both sides and with ATP. Finally, a loop (K47-Q52) N-terminal to the IF helix of Kir6.2 has close interaction with SUR1’s TMD0-intracellular loop 1 (ICL1), ICL2 and ICL3 (i.e. L0). Here, a compact network of interactions stabilizes the Kir6.2-CTD close to the membrane and also ATP binding. In the CTD-down conformation, the Kir6.2 pre-IF helix loop becomes more distant from the SUR1-ICLs such that the Kir6.2-CTD is no longer tethered close to the membrane, which also impacts the ATP binding pocket (see below).

### ATP binding pocket

Rapid and reversible closure upon non-hydrolytic binding of ATP at Kir6.2 is a cardinal feature of K_ATP_ channels (Nichols et al., 1996). The improved map quality in the current study allowed us to refine interpretation of the ATP cryoEM density and the interaction network that coordinates ATP binding and follow how the ATP binding pocket differs in different conformations.

In our improved maps, the ATP density could be modeled with ATP in two alternative poses. In the first, the γ-phosphate is oriented upward towards R50 of Kir6.2, which is consistent with functional studies indicating that R50 interacts with the γ-phosphate of ATP (Trapp et al., 2003). This pose was used to model ATP density in our previously published structure bound to GBC and ATP (PDB: 6BAA) (Martin et al., 2017a) and also ATPγS bound to a rodent SUR1-39aa-Kir6.2 fusion K_ATP_ channel (Wu et al., 2018). In the second pose, the ATP’s γ-phosphate is oriented downward facing N335, which is also supported by functional data showing that N335Q decreases ATP sensitivity (Drain et al., 1998) and used to model ATP density bound to Kir6.2 in cryoEM structures of a human SUR1-6aa-Kir6.2 fusion K_ATP_ channel (PDB: 6C3O and 6C3P) (Lee et al., 2017). Of note, in our previously published GBC/ATP map, an unassigned protruding density in ATP was also observed, which we speculated to be contaminating Mg2+ (Martin et al., 2017a) but which can be well modeled by the alternative pose of the γ-phosphate. Thus, the cryoEM density of ATP we observed is likely an ensemble of the two possible γ-phosphate poses.

The improved map also showed clear cryoEM density for the side chain of K205 in the L0 of SUR1. We have previously proposed that K205 participates in ATP binding (Martin et al., 2017b) based on early finding that K205E reduces ATP inhibition (Pratt et al., 2012). However, our previously published cryoEM map (EMD-7073) did not show clear side chain density of K205 to allow definitive conclusion. In our current map, K205 side chain is clearly oriented to the bound ATP (Fig.4B), stabilizing interactions with the β- and γ-phosphates of ATP. Similar observations have been reported by Ding et al. (Ding et al., 2019). The role of K205 in ATP binding is further substantiated by Usher et al. in which binding affinity of a fluorescent ATP analogue to the channel was assessed by FRET measurements between the ATP analogue and a fluorescent unnatural amino acid ANAP (3-(6-acetylnaphthalen-2-ylamino)−2-aminopropanoic acid) engineered at Kir6.2 amino acid position 311. The study found that SUR1-K205A and K205E mutations reduce ATP binding affinity by ∼5 and 10-fold (Usher et al., 2020).

**Figure 4.**
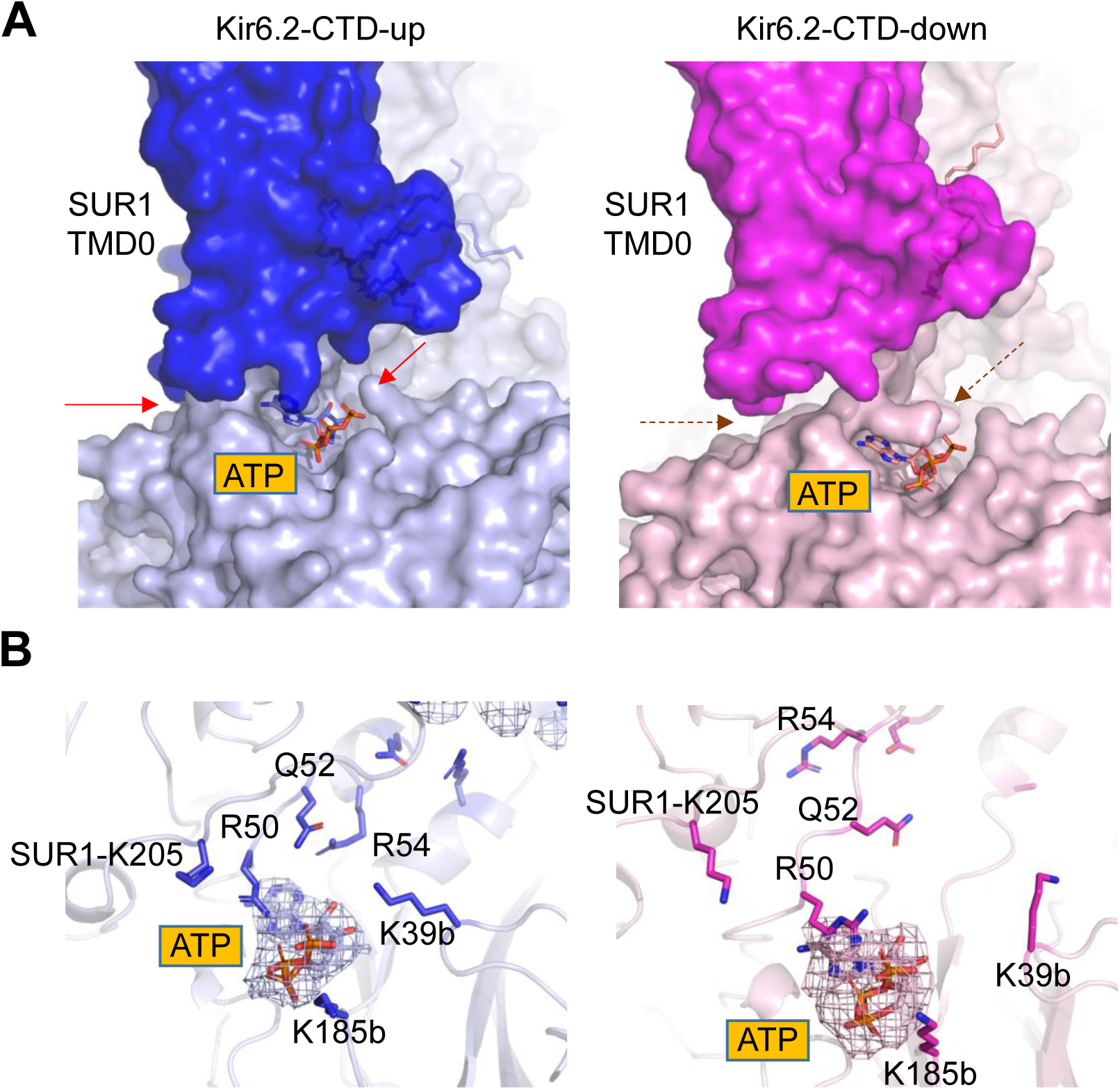
Comparison of he ATP binding pocket in Kir6.2-CTD-up and CTD-down conformations. **(A)** Comparison of the SUR1-Kir6.2 interface near the ATP binding pocket in Kir6.2-CTD-up (left) and the CTD-down (right) conformations in space filling model. The dashed arrows in the CTD-down panel point to contacts between SUR1 and Kir6.2 in CTD-up panel that are lost. The loss of the tight interaction renders ATP binding pocket less compact. **(B)** Residues surrounding ATP in the Kir6.2 CTD-up conformation (left) which showed changes in distance from ATP and/or side chain orientation in the CTD-down conformation (right), including K205 of SUR1, Q52, R54 and K39 of Kir6.2. ATP cryoEM density is represented by the mesh.

The conformational dynamics in Kir6.2-CTD and SUR1 have significant impact on the ATP binding site. In the Kir6.2-CTD-up conformation, the Kir6.2-CTD is packed tightly against SUR1’s ICL1, ICL2, and the initial segment of L0, and ATP fits snugly in the pocket formed by the N- and C-terminal domains of adjacent Kir6.2 subunits and L0 of SUR1. As Kir6.2-CTD rotates away towards the cytoplasm to the CTD-down conformation, it becomes disengaged from the SUR1-ICLs (Fig.4A), which disrupts several interactions that stabilize ATP binding. Specifically, R54, which is oriented towards the ATP in the CTD-up state becomes distant from the ATP binding pocket in the CTD-down structure. K39 in the N-terminus of neighboring Kir6.2, which also coordinates ATP binding in the CTD-up structure, is reoriented away from the ATP in the CTD-down structure. Moreover, the distance between K205 in the L0 of SUR1 and ATP is increased in the CTD-down conformation. These changes weaken the ATP binding pocket such that ATP becomes partially exposed to solvent. In addition Kir6.2-CTD dynamics, SUR1 rotation also affects ATP binding. When SUR1-ABC module rotates toward the Kir6.2 tetramer core (SUR1-in), L0 is pulled away from the ATP binding pocket. As such, SUR1-K205 loses interaction with ATP to destabilize ATP binding (Fig.4B).

### PIP_2_ binding site

At the PIP_2_ binding pocket, a lipid cryoEM density is seen in both Kir6.2-CTD-up and CTD-down conformations. Interestingly, the lipid density in the CTD-up conformation is significantly larger than that in the CTD-down conformation (Fig.5A). This was consistent for all datasets that have both conformations. We were able to fit, and tentatively model, the lipid density in the CTD-up structure with two PS molecules and that in the CTD-down structure with one PS molecule (Fig.S4). Since no PIP_2_ was added to our samples prior to imaging, we reasoned that the more abundant PS may have entered the binding pocket. In a recent study by Zhao and MacKinnon (Zhao and MacKinnon, 2021), it was shown that PIP_2_ is not required for K_ATP_ channel activity, suggesting other phospholipids that occupy the PIP_2_ binding site could potentially support channel activity. Whether the density in our structure represents PS, co-purified endogenous PIP_2,_ or other phospholipids requires further investigation.

**Figure 5.**
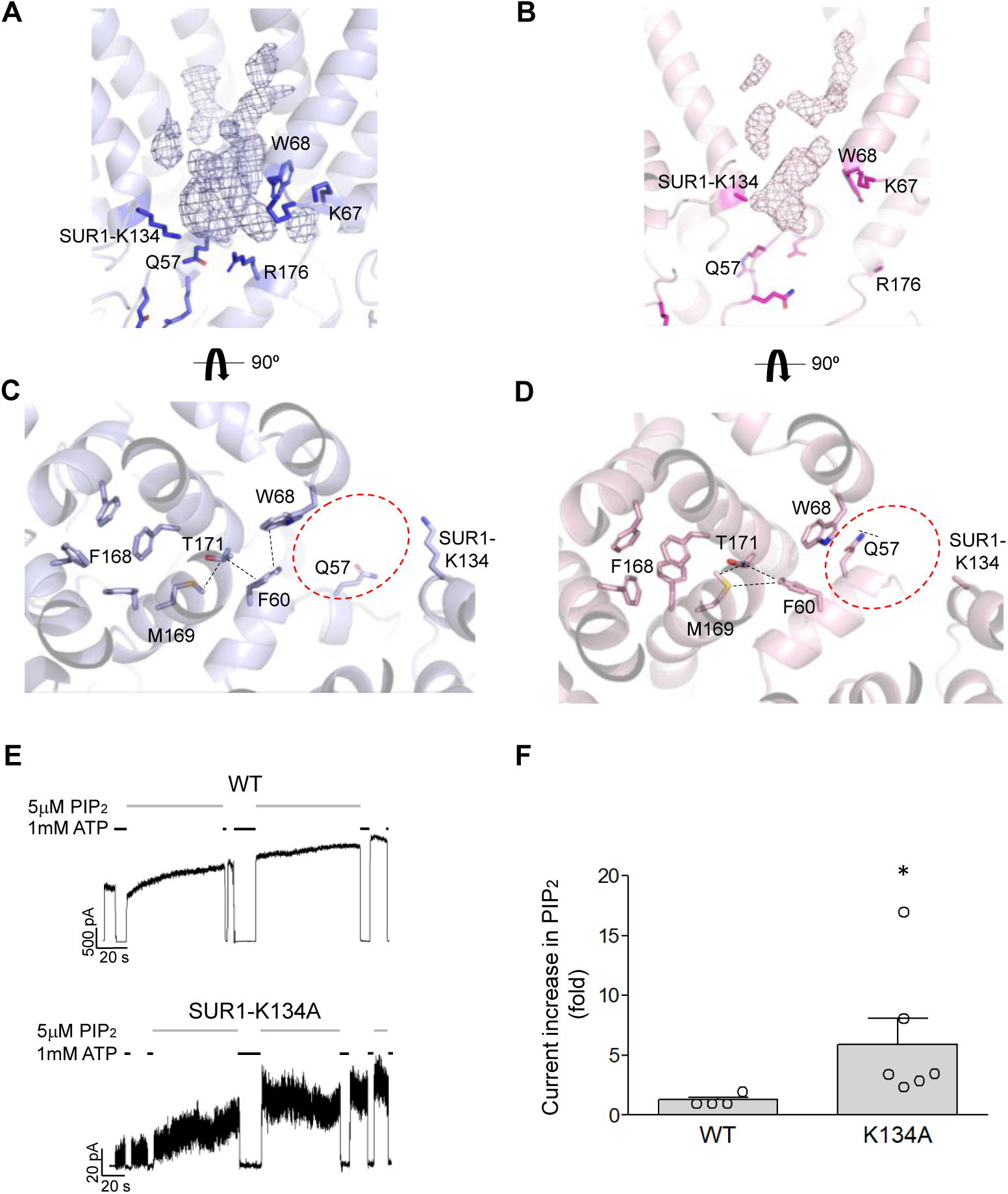
Comparison of the PIP_2_ binding pocket in Kir6.2-CTD-up and CTD-down conformations. **(A, B)** PIP_2_ binding pocket of Kir6.2-CTD up (A) and Kir6.2-CTD down (B) conformations viewed from the side. Lipid densities are shown in mesh. In (A), in addition to Kir6.2 residues previously implicated in phospholipid binding, SUR1-K134 side chain is pointed directly at the lipid density. **(C, D)** Same as (A, B) viewed from the extracellular side. The PIP_2_ binding pocket in both conformations are marked with broken circle with lipids removed. Note the PIP_2_ binding pocket is significantly more compressed in the Kir6.2-CTD-down than the CTD-up conformation due to secondary structural changes at the IF helix (from a 3_10_ helix to a straight shorter helix). **(E, F)** Inside-out patch-clamp recording examples showing greater fold current increase in response to PIP_2_ of the SUR1-K134A mutant channel than WT channel (left), with statistically significant difference (\**p*<0.05, student’s t-test).

In the Kir6.2-CTD-up structure, in addition to Kir6.2 K67, W68, and R176 previously implicated in PIP_2_ binding (Brundl et al., 2021; Cukras et al., 2002; Shyng and Nichols, 1998), K134 in TMD0 of SUR1 comes into close contact with the density corresponding to lipid headgroups (Fig.5A). To test whether this residue has a functional role, we mutated it to alanine and assessed PIP_2_ sensitivity. Compared to WT channels, SUR1-K134A mutant exhibited smaller initial currents in ATP-free solution in inside-out patch-clamp recording (see Methods). Upon PIP_2_ addition, currents increased by 5.93±2.17-fold, which is significantly higher than the 1.27±0.23-fold current increase seen in WT channels (Fig.5E, F), indicating the SUR1-K134A mutant reduced intrinsic *P*_*o*_ and PIP_2_ interactions. It is well documented that the Kir6.2 channel itself has low PIP_2_ sensitivty and low *P*_*o*_, but co-expression with SUR1 or just the TMD0 domain of SUR1 increases channel sensitivity to PIP_2_ and channel *P*_*o*_ by more than 10-fold (Babenko and Bryan, 2003; Chan et al., 2003; Enkvetchakul et al., 2000; Pratt et al., 2011). Our structure and function results suggest SUR1-TMD0 participates in PIP_2_ interaction, at least in part through K134, and strengthens PIP_2_ interactions with Kir6.2.

In the CTD-down structure, the IF helix is closer to W68 in the outer helix of the neighboring Kir6.2, causing compression of the PIP_2_ binding pocket (Fig.5A-D). This may explain why the lipid cryoEM density in the CTD-down structure is significantly smaller than that in the CTD-up structure and can be tentatively fit by only one PS molecule (Fig.S4). Moreover, because of the unwinding of the C-linker the key PIP_2_-interacting residue R176 is too distant for interaction. In this conformation, Kir6.2 is expected to be inactive.

### Elucidating the relationship between ATP and PIP_2_ binding by MD simulations

ATP and PIP_2_ compete with each other to close and open the channel, respectively. However, the structural mechanism underlying this functional competition remains unresolved. Mutation-function correlation studies have previously led to a proposal that ATP and PIP_2_ have overlapping but non-identical binding residues (Cukras et al., 2002; Shyng et al., 2000; Tucker et al., 1998). To test this hypothesis, we conducted MD simulation studies using the Kir6.2 (32-352) plus SUR1-TMD0 (1-193) tetramer part the RPG/ATP Kir6.2-CTD-up structure as a starting point. Previous studies have shown that Kir6.2 and TMD0 of SUR1 form “mini K_ATP_ channels” (Babenko and Bryan, 2003; Chan et al., 2003), which like WT channels exhibit functional antagonism between ATP and PIP_2_ (Babenko and Bryan, 2003; Pratt et al., 2011). The mini K_ATP_ channel system is therefore suitable for simulating residues which may participate in binding of both ligands. Three independent 1 μs simulations were carried out without ATP or PIP_2_ (apo) or with both ATP and PIP_2_ in their respective binding pockets (ATP+PIP_2_) (Fig.S5A, B; for details see Methods).

Comparing the two different conditions, there was an overall increased dynamics of the Kir6.2-CTD in the apo simulations versus the ATP+PIP_2_ simulations (Fig.6, Fig.S5). First, in the apo simulations, significant secondary structural changes at the IF helix and the C-linker were observed, resembling changes from the CTD-up to the CTD-down conformation observed in cryoEM structures. Second, the entire CTD appeared to relax towards the cytoplasm in the apo simulations. This was quantified by measuring the distance between the helix bundle crossing (HBC) gate at F168 and the G-loop gate at G295 (Fig.6A). In apo simulations, this distance increased over time in all three runs, whereas it remained relatively unchanged for the ATP+PIP_2_ simulations (Fig. 6B), except in run 2 during which the distance increased when ATP became partially dissociated at around 500 ns (Fig.6B). These findings show that in the absence of ligands, the Kir6.2-CTD has a tendency to relax into the CTD-down conformation.

**Figure 6.**
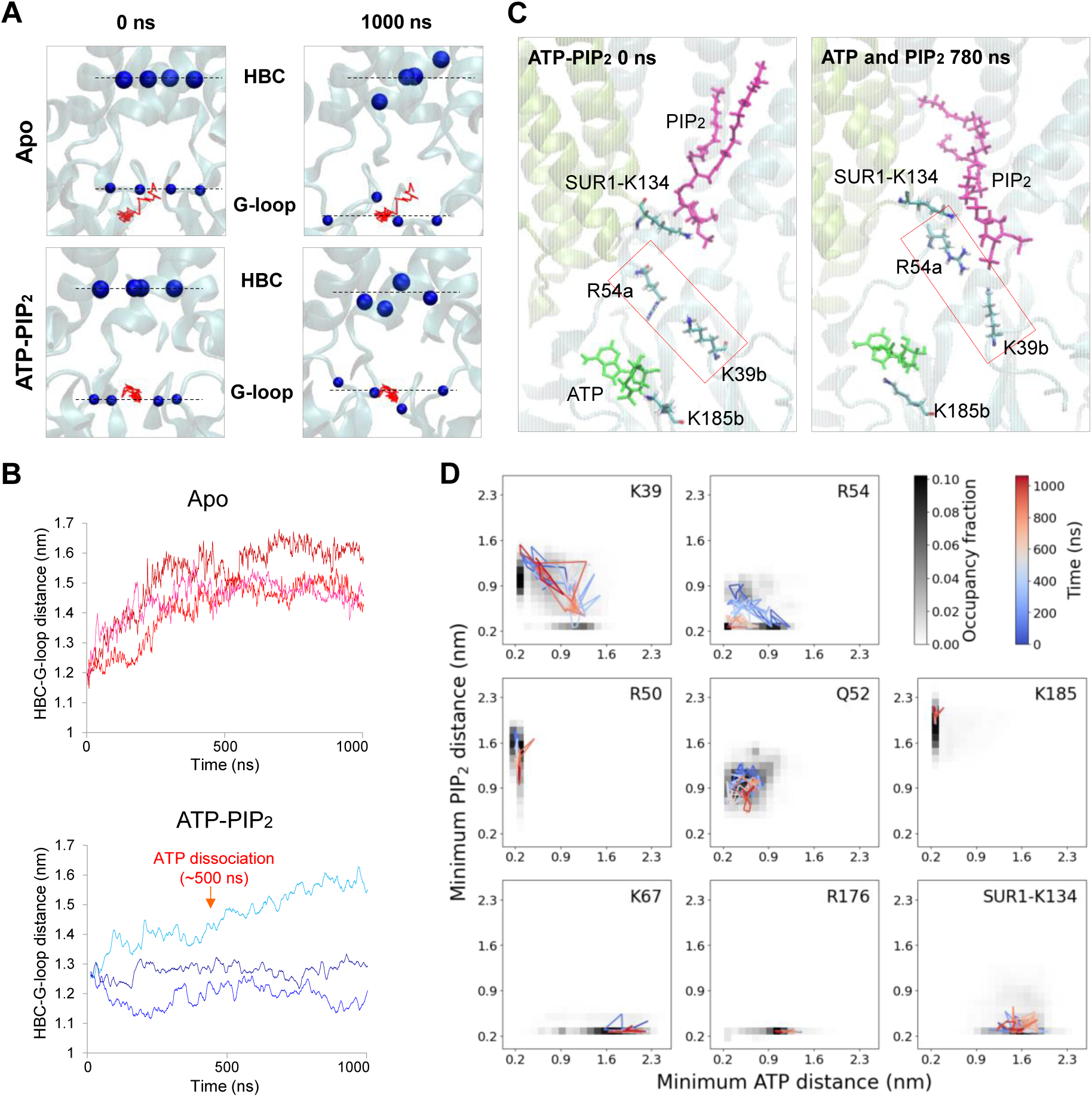
MD simulations of Apo versus ATP+PIP_2_ state. **(A)** Positioning of the helix bundle crossing (HBC) and G-loop gate in one apo-simulation and one ATP-PIP_2_-simulation. After aligning trajectories to the TM domain of the Kir6.2 channel, the time-varying position of the geometric center of G295 (G-loop gate) Cα carbons of all four chains (red) is overlaid on beginning and end snapshots of G295 Cα carbons. **(B)** Plots of the distance between the geometric centers of all four Cα atoms of F168 and of G295 for the entire simulations. The greater distances between the two gates in the apo state compared to the ATP+PIP_2_ state indicate relaxation of the CTD in the absence of ligands. In one apo simulation (light blue), ATP became partially dissociated at around 500 ns (red arrow). **(C)** Snapshots of simulations in the presence of ATP and PIP_2_ showing interactions of R54 and K39 with either ATP or PIP2. Kir6.2-K185 and SUR1-K134 which only interact with ATP or PIP_2_ respectively are also shown. **(D)** Heatmaps showing fraction of time residues spend at various distances from ATP+PIP_2_. A representative trajectory corresponding to chain 1 in run 1 and colored to show time evolution is shown for each residue. K39 and R54 (top row) exhibit switching behavior and occasional simultaneous contact, while essentially exclusive ATP binding residues are shown in the middle row, and PIP_2_ binding residues in the bottom row.

Analysis of the minimum distance between either ligands and their surrounding residues within 4 Å over the entire simulation shows that K39 and R54 of Kir6.2 engaged in both ATP and PIP_2_ binding. Fig.6D shows the fraction of time over the entire simulation each residue in each subunit and each run came into contact with ATP or PIP_2_. R54 and K39 each showed partial ATP and PIP_2_ occupancy (Fig.6C,D; Fig.S5C,D), which was in contrast to well established ATP binding residues, such as R50 and K185, or PIP_2_ binding residues such as K67 and R176, which showed nearly 100% ATP or PIP_2_ occupancy. Of the two, R54 showed greater interactions with both ATP and especially with PIP_2_, compared to K39. K39, while showing interaction with ATP in all three runs, only showed significant interaction with PIP_2_ in one of the three runs (Fig.S5D). The analysis also identified residues that had specific, although not completely stable, interactions with either ATP or PIP_2_ as defined by distance between residue and ligand <4 Å. In particular, Kir6.2-Q52 specifically interacted with ATP, and SUR1-K134 with PIP_2_, contrasting with the dual ligand binding mode of R54 and K39. Previous mutagenesis study has shown that mutation of K39 or R54 to alanine reduces channel open probability as well as sensitivity to ATP inhibition (Cukras et al., 2002), implicating a role of these residues in channel gating by PIP_2_ and ATP. Moreover, mutations R54C and R54H have been linked to congenital hyperinsulinism (De Franco et al., 2020) and mutation K39R to transient neonatal diabetes (Zhang et al., 2015). Our MD simulation results suggest both residues participate directly in ATP and PIP_2_ binding, providing mechanistic insight into how mutation of these residues affect PIP_2_ and ATP sensitivities and cause disease.

## Discussion

Cryo-preserved purified protein samples may contain multiple protein structures representing distinct functional or transitional states to provide mechanistic insight (Nogales and Scheres, 2015). In this study, analysis of five K_ATP_ channel cyroEM datasets collected in different ligand conditions revealed conformational heterogeneity of the Kir6.2-CTD and SUR1 ABC module. We observed the Kir6.2-CTD in either an “up” position tethered close to the plasma membrane, or a “down” position corkscrewed away from the membrane towards the cytoplasm. The ratio of the two conformations correlated with occupancy of inhibitory ligands at the SUR1 and Kir6.2 binding sites (Fig.1), suggesting inhibitory ligands help stabilize the Kir6.2-CTD close to the membrane. Furthermore, in both Kir6.2-CTD conformations the SUR1 ABC module was observed oriented with a range of rotation around the Kir6.2 tetramer central axis (Fig.S3). We observed a restructuring of protein-protein and protein-ligand interfaces in different conformations that sheds light on how ligands shift channel conformational dynamics to regulate gating.

### Correlation between Kir6.2-CTD conformation and channel function

The structures analyzed in this study all represent closed channels. Recently, an open human K_ATP_ channel structure containing Kir6.2 C166S and G334D mutations was reported (Zhao and MacKinnon, 2021), which showed a Kir6.2-CTD that is further CW rotated (extracellular view) and slightly upward translated compared to our Kir6.2-CTD-up conformation (Fig.1C). Rearrangement of the molecular interactions between the IF helix, the C-linker, and TM residues as well as increased distance between the pre-IF helix loop and SUR1 L0 compared to the ATP-bound closed WT channel structure (equivalent to our Kir6.2-CTD-up structure) were observed. The restructuring widens the ATP-binding pocket, explaining the absence of ATP cryoEM density despite high concentrations of ATP in the sample, and stabilizes HBC gate opening via side chain interactions between F60 in the IF helix and the HBC gate residue F168. Taken together, a picture emerges where the rotational and translational position of the Kir6.2-CTD determines the functional state of the channel. When the CTD is in the down position, the phospholipid binding pocket is compressed due to a secondary structural change at the IF helix (Fig.5), and the C-linker is unwound such that the CTD is distant from the membrane. We propose this conformation corresponds to an “inactivated” state in which the CTD is unable to engage with membrane phospholipids and thus open the channel. When the CTD is in the CW up-screwed position with ATP and/or drug bound, the channel is arrested in an inhibited state due to an interaction network between SUR1-L0 and Kir6.2-N terminal domain, which is stabilized by ATP and prevents further rotation of the CTD needed to open the HBC gate. Finally, when the CTD rotates further CW upon ATP dissociation, the gate widens to open the channel.

We propose the Kir6.2-CTD undergoes dynamic transitions between three conformations: CTD-down inactivated, CTD-up inhibited, and CTD-up open, and the probability of it in a given conformation is driven by ligands. In the absence of ligands the CTD-down conformation dominates, as seen in our apo state dataset, and the channel is inactivated. Binding of physiological inhibitor ATP at Kir6.2, and/or a pharmacological inhibitor at SUR1, shifts Kir6.2 towards the CTD-up but inhibited conformation. Binding of phospholipids, when coupled with unbinding of inhibitory ligands, shifts the equilibrium towards the Kir6.2-CTD-up open position. Under physiological conditions with high intracellular ATP concentrations and ambient PIP_2_, we envision the Kir6.2-CTD to be mostly in the ATP inhibited CTD-up conformation, with a small fraction in the phospholipids bound CTD-up further-rotated-open conformation; the CTD-down inactivated conformation would be rare. However, in pathological conditions the CTD-down inactivated state could be prevalent. We have previously reported several disease mutations at the Kir6.2 subunit-subunit interface (such as E229A, R314A, R192A, E227A) that cause channel inactivation (Lin et al., 2003). In channels containing such mutations, channels briefly open upon patch excision into ATP-free solution but then quickly inactivate. Interestingly, inactivation can be overcome by exposing channels to high concentrations of ATP followed by washout of ATP. We propose that these disease mutations increase the energy barrier for Kir6.2-CTD to transition from the CTD-down conformation to CTD-up conformation, thus trapping the CTD in the down inactivated state. ATP effectively acts as a molecular glue at its Kir6.2 domain interfaces. Exposure to ATP shifts the Kir6.2-CTD back to the CTD-up position such that channels can open again when ATP is washed out. The model similarly explains the ability of PIP_2_ to prevent and reverse inactivation mutants from inactivation by stabilizing Kir6.2-CTD in the up and further rotated open position.

The Kir6.2-CTD-up and CTD-down conformations observed in our structures are similar to the T and R states observed in a SUR1-39aa-Kir6.2 fusion channel bound to GBC+ ATPγS or ATPγS alone by Wu et al. (Wu et al., 2018). However, the percentage of particles in the T-state corresponding to our Kir6.2-CTD-up state in their ATPγS+GBC or ATPγS datasets is ∼40% and 43% respectively, which differ significantly from the ∼93% and 22% in our GBC+ATP and ATP alone datasets. The higher percentage of T-state particles in their ATPγS condition than that of CTD-up state in our ATP condition could be due to the 10-fold higher concentrations of ATPγS used for their structure but the relatively low percentage of T-state particles in their GBC+ATPγS condition is difficult to explain. We speculate the extra 39aa linker between SUR1 C-terminus and Kir6.2 N-terminus in the fusion construct may somehow uncouple drug binding from Kir6.2-CTD conformation. Consistent with this, GBC was shown to be ineffective in inhibiting the fusion channel in contrast to the WT channel formed by separate SUR1 and Kir6.2 proteins (Wu et al., 2018).

Of all pancreatic K_ATP_ channel cryoEM structures reported to date, only those formed by separate SUR1 and Kir6.2 proteins without SUR1 NBD dimerization show cryoEM density of the KNtp (Martin et al., 2019). The structures presented here refined our view of the molecular interactions between the KNtp and different parts of SUR1. We have previously shown that engineered crosslinking between Kir6.2-L2C and SUR1-C1142 reduces channel activity (Martin et al, 2019). A likely scenario is that stapling KNtp along SUR1 via the contact sites we observe stabilizes the inhibited Kir6.2-CTD-up conformation and prevents further rotation of the CTD needed to open the channel. This explains why deletion of KNtp increases channel open probability (Babenko et al., 1999; Koster et al., 1999; Reimann et al., 1999), while drugs which stabilize KNtp in the transmembrane cavity of the SUR1 ABC module such as GBC, RPG and CBZ mimic the physiological inhibitor ATP and block channel activity (Devaraneni et al., 2015).

### Comparison to other Kir channels

Differential CTD rotation and proximity to the membrane have also been reported for other Kir channels. Crystal structures of Kir2 have shown that PIP_2_ and other anionic phospholipids anchor the CTD close to the membrane to allow channel opening (Lee et al., 2016b; Zangerl-Plessl et al., 2020). A recent cryoEM study of Kir3 channels found that without PIP_2_, the CTD is CCW rotated away from the membrane and increasing PIP_2_ shifts the CTD to a CW rotated (extracellular view) and upward translated position close to the membrane (Niu et al., 2020), which are qualitatively similar to the Kir6.2 CTD-down and CTD-up conformations presented here. These findings indicate a common theme in Kir channel conformation and gating transitions. However, unlike Kir2 and Kir3 channels, K_ATP_ channels have an additional ATP-bound Kir6.2-CTD-up conformation stop between the CTD-down inactivated and CTD-up open conformations to allow rapid and reversible inhibition of the channel in response to metabolic signals.

Another unique feature of Kir6.2 channels is the requirement of SUR1 co-assembly to achieve the high ATP and open probability of native K_ATP_ channels (Inagaki et al., 1995; Tucker et al., 1997). Several studies have now provided functional, biochemical and structural evidence that SUR1 directly participates in ATP binding via K205 in the L0 linker (Ding et al., 2019; Pratt et al., 2012; Usher et al., 2020). In addition, our structures suggest SUR1 via its TMD0 ICLs and L0 also form a network interaction with Kir6.2 to stabilize it in the CTD-up position to enhance ATP sensitivity. SUR1 increases the open probability of Kir6.2 by more than 10-fold, an effect that is largely mediated by TMD0 (Babenko and Bryan, 2003; Chan et al., 2003). We show in our structure that K134 in SUR1-ICL2 is oriented towards the lipid headgroup density in the PIP_2_ binding pocket, suggesting SUR1-TMD0 increases channel open probability by directly contributing to binding of PIP_2_ or other phospholipids. Supporting this, MD simulations show high PIP_2_ occupancy (Fig.6D, Fig.S5D) and functional experiments show that mutation of SUR1 K134 to alanine reduces channel P_o_ (Fig.5E, F).

### Structural basis of ATP and PIP_2_ antagonism

ATP and PIP_2_ functionally compete to inhibit and activate K_ATP_ channels, respectively (Baukrowitz et al., 1998; Shyng and Nichols, 1998). Molecular dynamics (MD) simulations reveal two Kir6.2 residues, K39 and R54, interact with both ATP and PIP_2_, providing evidence that competition for overlapping binding residues between the two ligands underlies, at least in part, functional competition between ATP and PIP_2_. Interestingly, in one of the ATP+PIP_2_ simulation runs, ATP dissociates from its binding pocket (Fig.6B). This dissociation event is likely captured because SUR1-L0, which contains the ATP stabilizing residue K205 is not included in the simulation structure. It offers a hint of how ATP may dissociate, leaving binding residues shared between ATP and PIP_2_ greater freedom to interact with PIP_2_ to favor channel opening. The rotational movement of SUR1 towards the Kir6.2-tetramer observed in our multibody refinement analysis increases the distance between SUR1-K205 and bound ATP (Fig.S3), which may initiate ATP dissociation by weakening ATP binding thus providing a pathway for channel transition from ATP bound inhibited state to PIP_2_ bound open state.

In summary, the structural analysis, MD simulations, and functional studies presented here together with the recent open K_ATP_ channel structure reported by others offer insight into several longstanding questions on K_ATP_ channel gating mechanisms. A question which remains unresolved is the full extent of the conformational dynamics of the SUR1 subunit and how it relates to channel function. Large rotation of the SUR ABC module that leads to a quatrefoil channel conformation has previously been reported in a human SUR1-6aa-Kir6.2 fusion protein channel with MgATP/MgADP bound dimerized NBDs (Lee et al., 2017). Recently, a similar large rotation is reported in a SUR2B/Kir6.1 vascular K_ATP_ channel bound to GBC and ATP (Sung et al., 2021). However, the quatrefoil conformation was not observed in the most recent human SUR1 NBDs dimerized-Kir6.2 mutant open channel (Zhao and MacKinnon, 2021). Whether the variable findings stem from protein samples or due to differences in data processing will be important to resolve in order to fully understand K_ATP_ channel structure and function relationship.

## Methods

### Image processing and particle classification

CryoEM images of pancreatic K_ATP_ channels (co-assembled from hamster SUR1 and rat Kir6.2) collected in different liganded conditions in our previous publication (Martin et al., 2019) were reprocessed from the initial 2D classification step that we described previously using RELION-3.1 (Zivanov et al., 2018). Classes displaying fully and partially assembled complexes with high signal/noise were selected. The particles were re-extracted at 1.045 Å/pix for RPG/ATP, 1.399 Å/pix for GBC/ATP, 1.72 Å/pix for CBZ/ATP and ATP only, and 1.826 Å/pix for apo state datasets, and then used as input for 3D classification in RELION-3.1. Fig.S1A shows the data processing workflow for the RPG/ATP dataset. Channel particles refined in the final C4 reconstruction (150,707 total particles) were subjected to C4 symmetry expansion and yielded 4 fold more copies. Further refinement was performed without symmetry restraints or masking such that possible heterogeneous particles can be aligned without any restraints. 2D class averages from all data sets showed significant heterogeneity of the SUR1-ABC module, indicating dynamic SUR1 motions were captured during vitrification of the cryoEM samples. To probe potential novel conformations due to dynamics of SUR1-ABC module relative to the Kir6.2 tetramer, or for novel conformations that arise due to dynamic motions of other domains of the K_ATP_ channel, a soft mask that includes the Kir6.2 tetramer and one SUR1 in a propeller form was created in Chimera using our previously published model (PDB:6BAA) (Martin et al., 2017a), and extensive focused 3D classification was performed without particle alignment. This revealed two major classes with different Kir6.2-CTD conformations that are either anchored up towards the plasma membrane (CTD-up) or extended down further towards the cytoplasm (CTD-down).

Focused refinement of SUR1 was carried out after partial signal subtraction that removed signals outside the masked region, followed by further 3D classification without alignment at higher regularization T values (ranged from 6 to 20) and local refinement of signal subtracted particles (Scheres, 2016). Extensive 3D classification sorted out remaining minor groups of particles that did not align well with the propeller conformation but no other conformations emerged. The dominant class then underwent three iterations of CTF refinement and 3D refinement. To test whether some of the SUR1 particles adopt quatrefoil-like conformation reported previously, a mask that includes the Kir6.2 tetramer and one SUR1 was also created using a quatrefoil-like model of our previously published Kir6.1/SUR2B structure (PDB:7MJO) (Sung et al., 2021), which was used for classification following the same scheme described for the propeller form mask. A minor quatrefoil form at 7.1 Å overall resolution was identified from the RPG/ATP dataset (Fig.S1A). Final maps were subjected to Map-modification implemented in *Phenix* with two independent half maps and corresponding mask and model as input. They were then sharpened with model-based auto sharpening with the corresponding model using *Phenix*, a step that was iterated during model building.

The same workflow was used to process the other four datasets, GBC/ATP, CBZ/ATP, ATP only and apo states. With the exception of the apo dataset which yielded only the CTD-down conformation, all other datasets showed both Kir6.2-CTD classes similar to those identified in the RPG/ATP dataset but with varying ratios of the two conformations. Particle distributions and final map resolutions for all datasets are summarized in Table S1. Upon carrying out this further analysis, we noted the CBZ/ATP dataset previously reported to be collected in the presence of 10μM CBZ and 1mM ATP did not yield a map with clear ATP density at the Kir6.2 ATP binding site. Upon inspection of the ATP used it was discovered that the concentration had been mistakenly reported as 1mM rather than 0.1mM, which likely explained the lack of ATP cryoEM density.

### Multibody refinement

For Kir6.2 tetramer multibody refinement, particles pooled from both the RPG/ATP CTD-up and CTD-down classes and also each class separately were used (Nakane et al., 2018). To interrogate the Kir6.2 CTD movements relative to its TM domain, we masked out the SUR1 density and assigned the Kir6.2 TM domain (58-173) and CTD (174-352) as two separate rigid bodies (Fig.S2A). Principal component analysis showed that a dominant eigenvector accounted for 25.3% of the overall variance (Fig.S2B). Histograms of the amplitudes along this eigenvector shows a bimodal distribution with two peaks, indicating two conformational distinct populations differing in the distance between the CTD and the TM domain and rotation of the CTD as expected. Similar rotation and rocking motions of the CTD were also observed within the Kir6.2-CTD-up and Kir6.2-CTD-down class particles, where greater heterogeneity was seen in the CTD-down than the CTD-up population of particles (Fig.2SC, D), indicating increased mobility of the CTD when it is detached from the membrane.

For (Kir6.2)_4_-SUR1 multibody refinement, the map was divided into three bodies: body 1: Kir6.2-tetramer (E30-D352), body 2: ABC-core (Q211-V1578) of SUR1 plus KNtp (M1-E19) of Kir6.2, and body 3: TMD0 (M1-L210) of SUR1 (Fig.S3A). Multibody refinement was repeated with varying standard deviations of degrees on the rotation and pixels on the translation to rule out artifacts. Principal component analysis in the *relion_flex_analyse* program revealed that approximately 17.5% of the variance is explained by the dominant eigenvector 1 (Fig.S3B,C) corresponding to horizontal swinging motion of SUR1 (Nakane et al., 2018). We then used the program to generate two separate STAR files, each containing ∼35,000 particles with eigenvalues less than −5 or greater than +5 along eigenvector 1 (Fig.S3D). These two sets of particles were further refined using a soft mask, yielding two maps which we refer to as SUR1-out and SUR1-in with an overall resolution of 3.8 and 3.9 Å, respectively.

### Model building and refinement

The RPG/ATP dataset yielded the highest resolution maps and were used for modeling (Table S2). Initial models for the Kir6.2_4_-SUR1_1_ channel were obtained by docking Kir6.2-TMD (32-171) and Kir6.2-CTD (172-352) from our previously published model (PDB:6BAA) (Martin et al., 2017b), and TMD0/L0 (1-284), TMD1 (285-614), NBD1 (615-928), NBD1-TMD2-linker (992-999), TMD2 (1000-1319) and NBD2 (1320-1582) of SUR1 (PDB:6PZA)(Martin et al., 2019) into either the RPG-CTD-up or the RPG-CTD-down cryoEM density map by rigid-body fitting using Chimera’s ‘Fit in’ tool (Pettersen et al., 2004). Then each domain was combined using Chimera and served as a template model. We used Coot to manually build and edit residues 32-78 and residues 79-361 at the interface of two Kir6.2 subunits (Emsley et al., 2010), we then copied those changes to each of the other four Kir6.2 subunits. Further edits and refinements were done independently to each chain without strict NCS restraints. The models were then iteratively built and refined in Coot (Emsley et al., 2010) and Phenix (Afonine et al., 2018), with Ramachandran restraints, secondary structure restraints, and side-chain rotamer restraints. The N-terminal of Kir6.2 (residues 1-31, KNtp) had sufficient continuous density and the density was sufficiently clear to allow modeling of several key interactions with SUR1. However, accurate modeling of side chains was not possible for the entire KNtp. Similarly, the NBDs of SUR1 had weaker density than most of the reconstructed map, and particularly NBD2 has weak density and our model building relied heavily on restraints and prior models. In addition to modeling the protein, ATP was modeled at the inhibitory ATP-binding site on each of the four Kir6.2 subunits and the nucleotide binding site in NBD1 of SUR1 where there was sufficient cryoEM density. Phosphatidylserine, phosphatidylcholine, and phosphatidylethanolamine were modeled liberally into plausible lipid density, 17 lipids were modeled for the RPG-CTD-up model and 13 lipids modeled for the RPG-CTD-down model.

### MD simulations and analysis

All MD simulations were performed at all-atom resolution using AMBER 16 (Case et al., 2016) with GPU acceleration. Initial coordinates were developed from the RPG/ATP-CTD-up model including four Kir6.2 (32-352) and four SUR1-TMD0 (1-193) without the SUR1-ABC core. ATP in the cryoEM structure was removed for simulations in the apo condition. For simulations in the presence of ATP and PIP_2_, ATP from the cryoEM structure was kept and a PIP_2_ molecule (DMPI24, di-myristoyl-inositol-(4,5)-bisphosphate) was docked in the PIP_2_ binding pocket using Kir3.2-PIP_2_ structure (PDB ID: 6M84) as a template.

The simulation starting structures were protonated by the H++ webserver (http://biophysics.cs.vt.edu/H++) at pH 7 and inserted in a bilayer membrane composed of 1-palmitoyl-2-oleoyl-phosphatidylcholine (POPC) lipids and surrounded by an aqueous solution of 0.15 M KCl. The optimal protein orientations in the membrane were obtained from the OPM database (Lomize et al., 2012). All systems contain 650-680 POPC lipids and ∼87,000 water molecules, resulting in a total of ∼385,000 atoms. They were assembled using the CHARMM-GUI webserver (Jo et al., 2008; Lee et al., 2016a; Wu et al., 2014), which also generated all simulation input files.

The CHARMM36m protein (Huang et al., 2017) and CHARMM36 lipid (Klauda et al., 2010; Pastor and Mackerell, 2011) force field parameters were used with the TIP3P water model (Jorgensen, 1983). Langevin dynamics (Pastor, 1988) were applied to control the temperature at 300K with a damping coefficient of 1/ps. van der Waals (vdW) interactions were truncated via a force-based switching function with a switching distance of 10 A° and a cutoff distance of 12 A°. Short-range Coulomb interactions were cut off at 12 A°, long-range electrostatic interactions were calculated by the Particle-Mesh Ewald summation (Darden et al., 1993; Essmann et al., 1995). Bonds to hydrogen atoms were constrained using the SHAKE algorithm (Jean-Paul Ryckaert, 1977).

The atomic coordinates were first minimized for 5000 steps using the steepest-descent and conjugated gradient algorithms, followed by a ∼2 ns equilibration simulation phase, during which dihedral restraints on lipid and protein heavy atoms were gradually removed from 250 to 0 kcal/mol/Å2, the simulation time step was increased from 1 fs to 2 fs, and the simulation ensemble was switched from NVT to NPT. To keep the pressure at 1 bar, a semi-isotropic pressure coupling was applied that allows the z-axis to expand and contract independently from the x-y plane (Martyna, 1994). The simulations were then run for over 1 μs with a time step of 4 fs enabled by hydrogen mass repartitioning (Balusek et al., 2019; Hopkins et al., 2015).

### Analysis of ATP/PIP_2_ occupancy

ATP and PIP_2_ residue occupancies (Fig. 6D) were computed using the histogram of minimum hydrogen bond lengths between each residue and PIP_2_/ATP, to show the amount of time spent at different distances. Summarized residue occupancies (Fig. S5D) were calculated as the fraction of time each residue spent in contact with ATP/PIP_2_, where contact is defined as a minimum hydrogen bond length of below 4 A°. For both, minimum bond lengths were used regardless of which pairs formed the bond. A 10 ns window average was used to smooth the minimum bond length time series data.

### Functional studies

Point mutation SUR1-K134A was introduced into hamster SUR1 cDNA in pECE using the QuikChange site-directed mutagenesis kit (Stratagene). Mutation was confirmed by DNA sequencing. For electrophysiology, wild-type or mutant SUR1 cDNA and rat Kir6.2 in pcDNA1 along with cDNA for green fluorescent protein GFP (to facilitate identification of transfected cells) were co-transfected into COS cells using FuGENE®6, and plated onto glass coverslips 24 hours after transfection for recording, as described previously (Martin et al., 2019). Recording pipettes were pulled from non-heparinized Kimble glass (Fisher Scientific) on a horizontal puller (Sutter Instrument, Co., Novato, CA, USA). Electrode resistance was typically 1-2 MΩ when filled with K-INT solution containing 140 mM KCl, 10 mM K-HEPES, 1 mM K-EGTA, pH 7.3. ATP was added as the potassium salt. PI4,5P_2_ (Avanti Polar Lipids) was reconstituted in K-INT solution at 5μM and bath sonicated in ice water for 20 min before use. Inside-out patches of cells bathed in K-INT were voltage-clamped with an Axopatch 1D amplifier (Axon Inc., Foster City, CA). Exposure of membrane patches to ATP- or PIP_2_-containing K-INT bath solution was as specified in Fig.5 legend. All currents were measured at room temperature at a membrane potential of -50 mV (pipette voltage = +50 mV) and inward currents shown as upward deflections. Data were analyzed using pCLAMP10 software (Axon Instrument). Off-line analysis was performed using Microsoft Excel programs. Data were presented as mean ± standard error of the mean (s.e.m) and statistical analysis was performed using two-tailed student’s *t*-test, with *p*<0.05 considered statistically significant.

## Acknowledgements

We thank Zhongying Yang for technical support. The project was supported by the National Institutes of Health grant R01DK066485 (to S.-L. S.) and the National Science Foundation grant MCB 17158233 (to D.M.Z.).

## Author contributions

MWS performed image analysis, atomic modeling, MD simulation analysis, prepared figures and wrote the manuscript. CMD performed atomic modeling, analyzed data, prepared figures and wrote the manuscript. BM performed MD simulations. JDR analyzed MD simulation data, prepared figures and contributed to manuscript preparation. BLP contributed to discussion of the project and edited the manuscript. DMZ provided guidance to the design and analysis of MD simulation studies and edited the manuscript. SLS conceived the project, performed electrophysiology experiemnts, analyzed data, prepared figures and wrote the manuscript.

## Competing interests

The authors declare that they have no competing financial or non-financial interests with the contents of this article.

---

**Figure S1.**
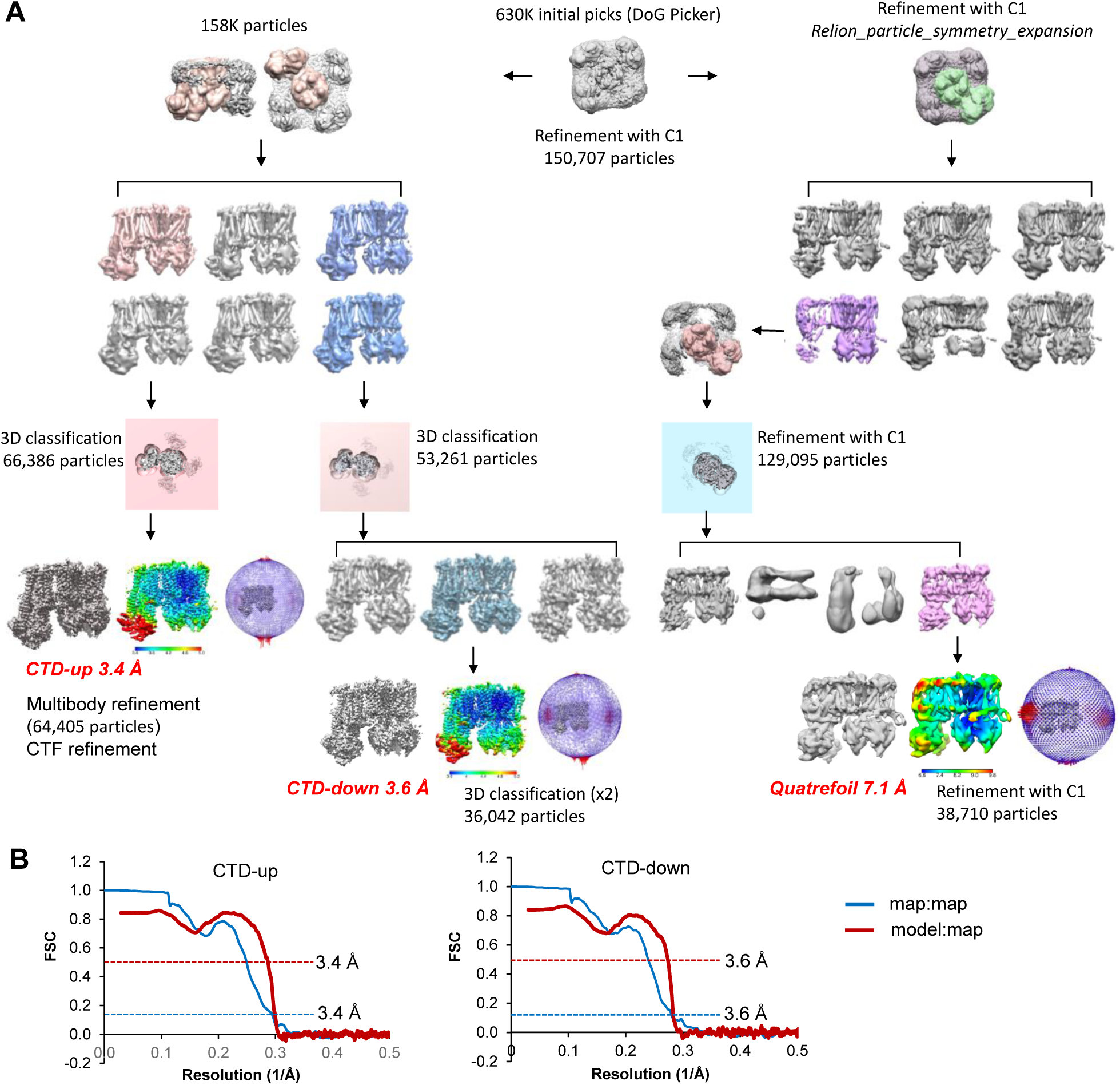
**(A)** Image data processing workflow for the RPG/ATP dataset. **(B)** Fourier Shell Correlation (FSC) curve between two independently refined half-maps (blue; resolution at FSC=0.143 shown on the right). The atomic model cross-validation curve (red) is the FSC between the map derived from the model and the postprocessed experimental full map. The resolution at FSC=0.5 is shown on the right.

**Figure S2.**
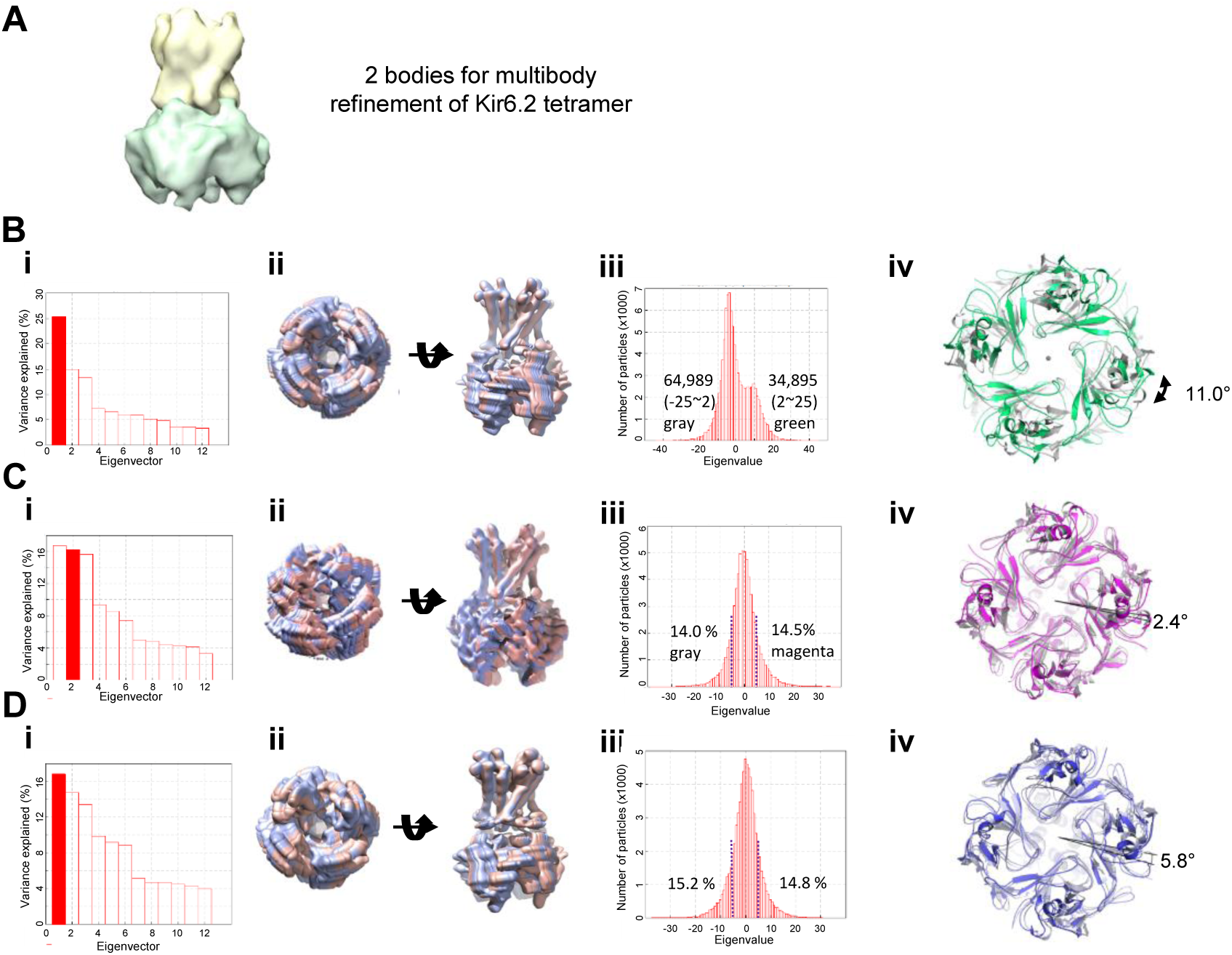
Analysis of Kir6.2 conformational dynamics by multibody refinement in Relion 3. **(A)** CryoEM maps of two classes (left:K4S-RPG-CTD-up and left:K4S-RPG-CTD-down) from 3D focused refinement of K4S showing the rotational and translational difference of the CTD in the RPG/ATP dataset. Kir6.2 domain including KNtp in each conformations is indicated in blue and magenta respectively, SUR1 domain is in faint pink. **(B)** Multibody refinement strategy to probe the Kir6.2-CTD movement relative to the TM domain. **(C)** Multibody refinement of Kir6.2 tetramer core using particles from both Kir6.2-CTD-up and CTD-down classes shown in (A). i. Eigenvectors vs variance plot; ii. Motion represented by the first eigenvector view from bottom and the side; iii particle distribution plot for eigenvector 1 showing bimodal distribution; iv, comparison of models refined in the maps from the first peak and the second peak showing rotation of the CTD viewed from the bottom. **(D)** The same analysis except only Kir6.2-CTD-up class of particles were included and eigenvector 2 representing CTD rotation movement is analyzed. (E) The same analysis except only Kir6.2-CTD-down class of particles were used. Eigenvector 1 motion was analyzed as it is the same rotational movement as eigenvector 2 in (D).

**Figure S3.**
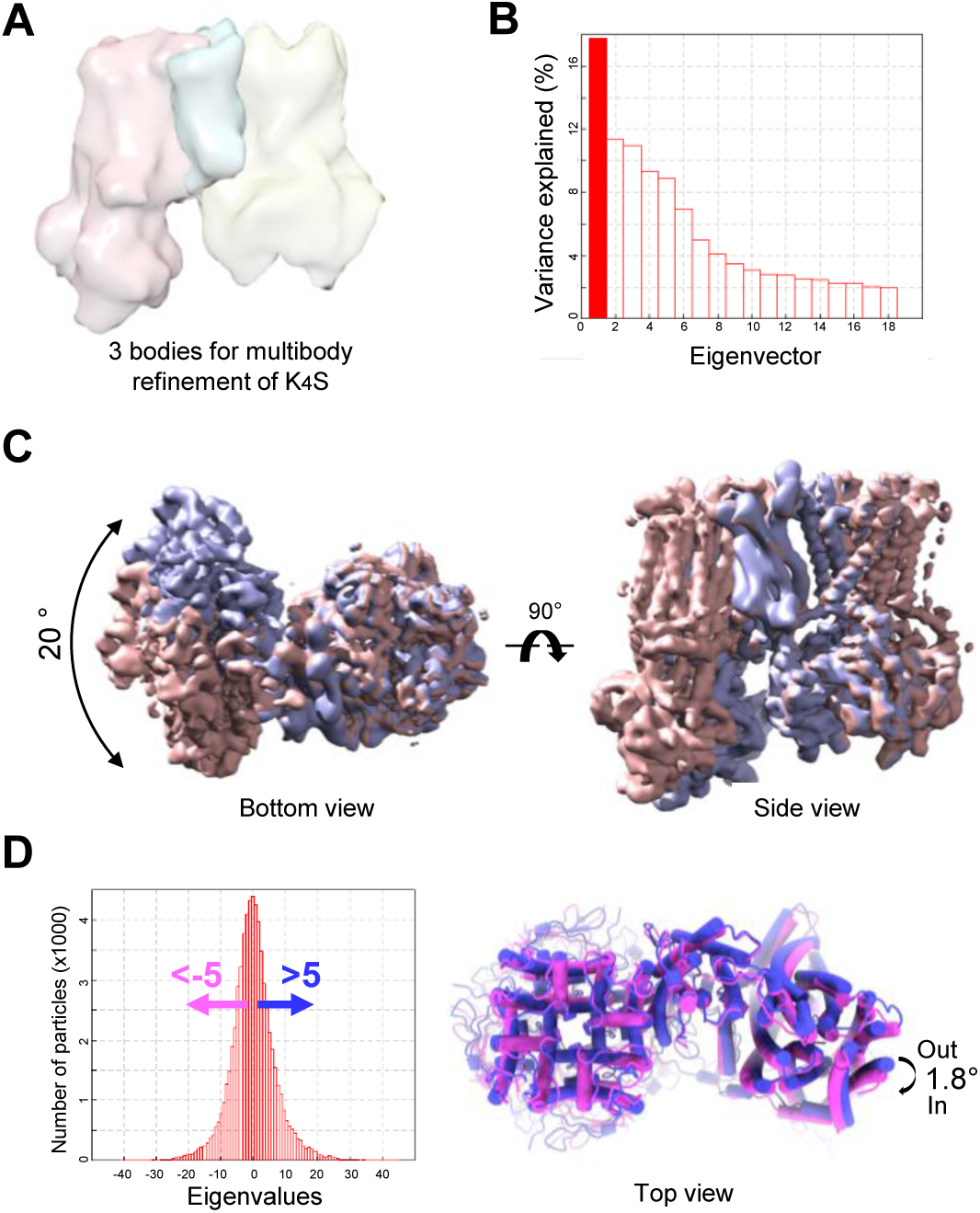
Analysis of SUR1 conformational dynamics by multibody refinement in Relion 3. **(A)** Multibody refinement strategy to probe the SUR1 movement relative to the TM domain using particles in the Kir6.2-CTD-up class from the K_4_S focused 3D classification. **(B)** Eigenvector vs variance plot. **(C)** Comparison of maps representing the extremes of the movement spectrum along eigenvector 1 with the swinging motion range of 20 degrees viewed from the bottom. Viewed from the side, the SUR1 ABC module is closer to Kir6.2 when it swings in towards Kir6.2 (lavender, SUR1-in conformation) compare to the conformation when SUR1 swings away from Kir6.2 (salmon, SUR1-out conformation). **(D)** Particle numbers vs eigenvalue along eigenvector 1. Particles with eigenvalues <-5 and >5 were separately refined for building SUR1-in (pink) and SUR1-out (blue) models, respectively.

**Figure S4.**
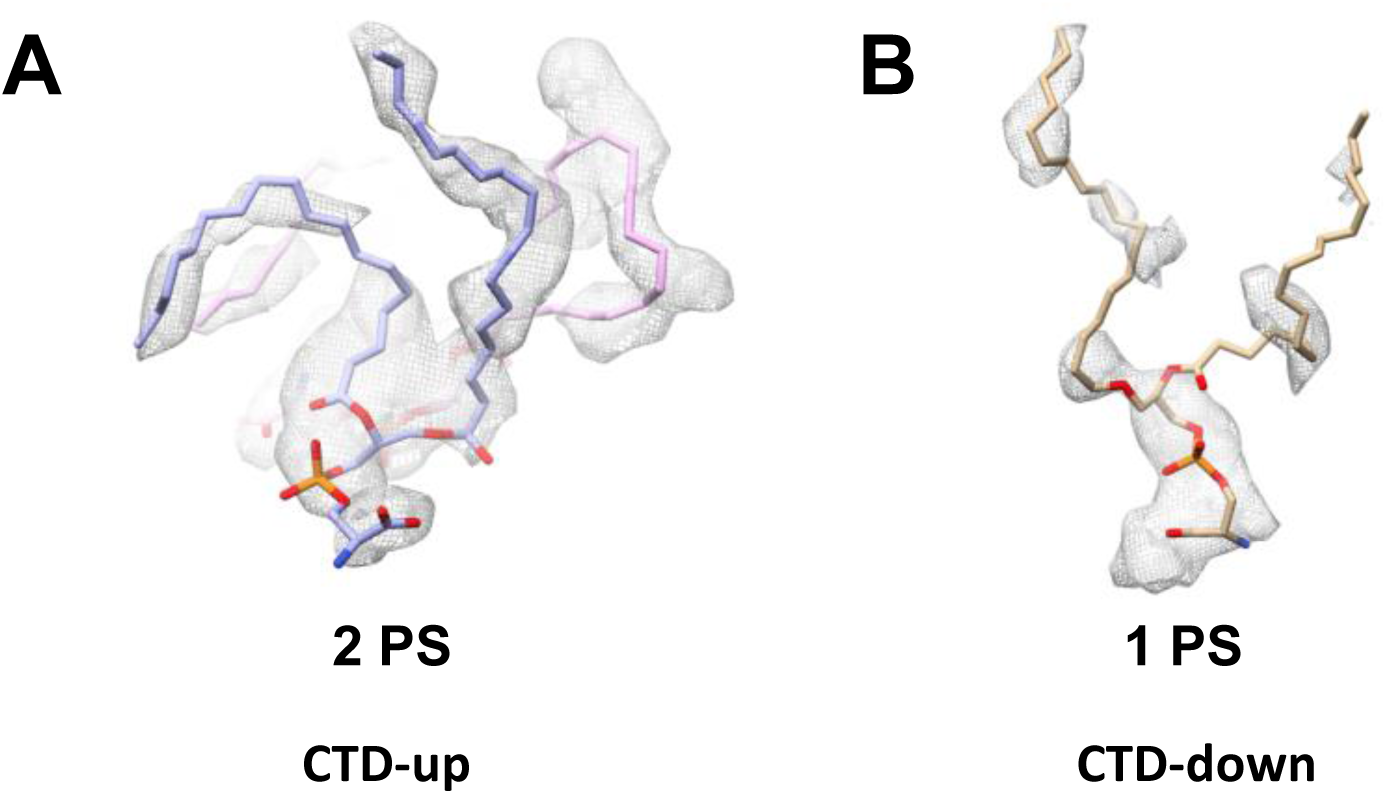
Lipid cryoEM density in RPG/ATP structures with **(A)** CTD-up structure tentatively fitted with two phosphatidylserine (PS) molecules and **(B)** CTD-down structure tentatively fitted with one PS molecule.

**Figure S5.**
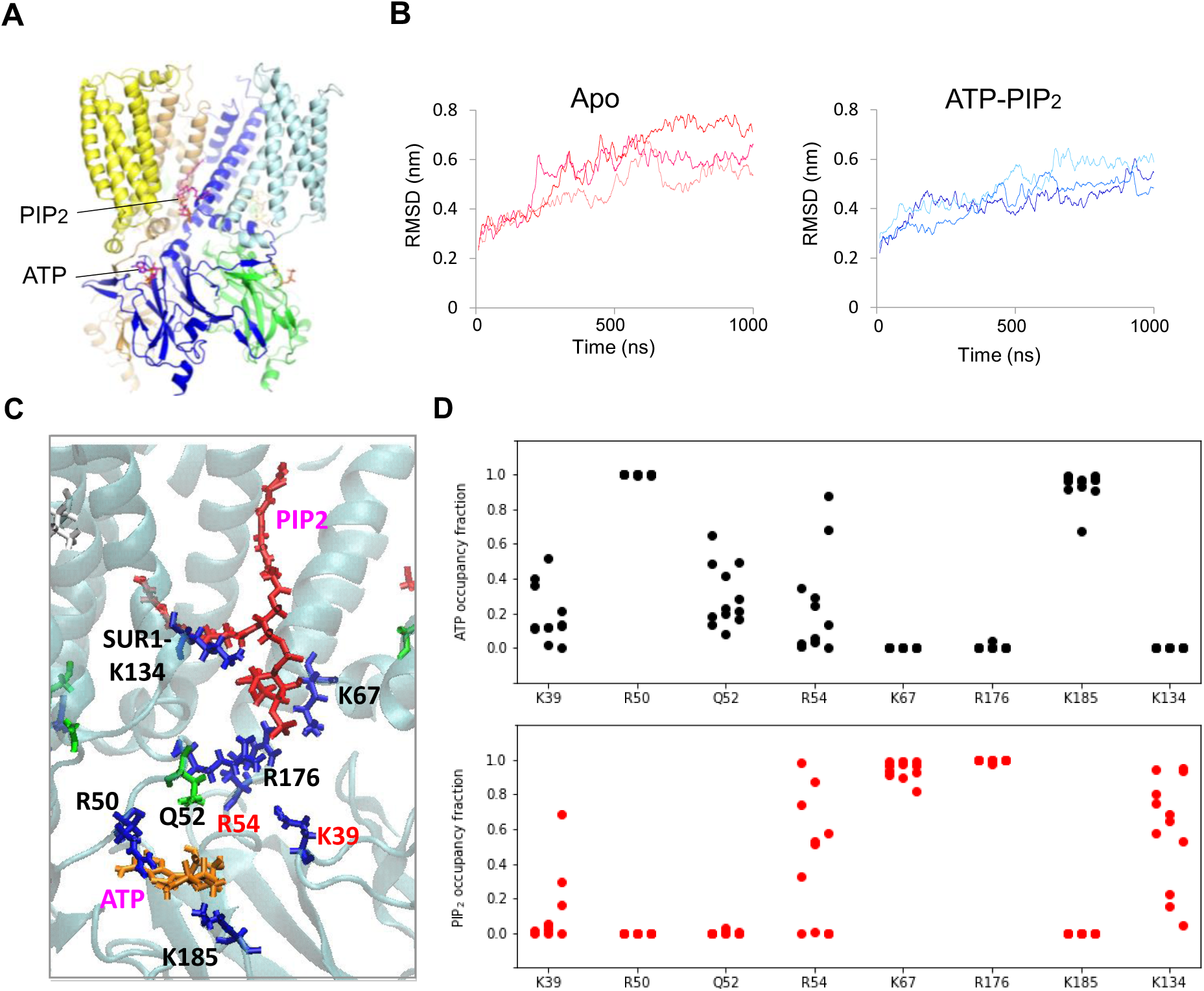
MD simulations. **(A)** Kir6.2 (32-352) and SUR1-TMD0 (1-196) model used for simulations. The starting model is from the RPG/ATP Kir6.2-CTD up conformation. For apo state simulation, bound ATP is removed. For ATP+PIP_2_ simulation, ATP is bound as in the structure and one PIP_2_ molecule is docked at the PIP_2_ binding pocket based on crystal structure of Kir3.2 in complex with PIP_2_. **(B)** RMSD plot for the three individual runs for apo (left) and ATP+PIP_2_ simulations. **(C)** Simulation snapshot showing residues included in the ATP and PIP2 occupancy analysis shown in (D). **(D)** Plots of fraction of time over the 1 μs simulation each residue on the x-axis came into within 4 Å (minimum distance of N-atoms of side chains and O-atoms of ligands) of ATP (ATP occupancy) or PIP_2_ (PIP_2_ occupancy). K185 of Kir6.2 and K134 of SUR1-TMD0 show relatively stable interactions with the oxygen of phosphate groups of ATP or PIP_2_ respectively; by contrast, R54 and K39 of Kir6.2 switch their interactions between ATP and PIP_2_.

**Table S1.** Summary of conformations observed in the five cryoEM datasets from K_4_S focused 3D classification.

|  |  | RPG/ATP | GBC/ATP | CBZ/ATP | ATP-only | Apo |
| --- | --- | --- | --- | --- | --- | --- |
| SUR1-focused (from Martin et al., 2019) | Resolution | 3.6 Å | 3.75 Å | 4.3 Å | 4.5 Å | 4.55 Å |
| Kir6.2-CTD-up | Resolution | 3.44 Å | 3.58 Å | 5.2 Å | 5.7 Å | N/A |
|  | Particle # (%) | 64,405 (64%) | 160,000 (92%) | 50,000 (22%) | 33,000 (18%) | - |
| Kir6.2-CTD-down | Resolution | 3.6 Å | 6.8 Å | 4.1 Å | 4.52 Å | 4.55 Å |
|  | Particle # (%) | 36,042 (36%) | 13,000 (8%) | 180,000 (78%) | 152,000 (82%) | 90,000(100%) |

**Table S2.** Model statistics.

|  | RPG/ATP | RPG/ATP | RPG/ATP, Kir6.2-CTD-up |  |
| --- | --- | --- | --- | --- |
|  | Kir6.2-CTD-up | Kir6.2-CTD-down | SUR1-in | SUR1-out |
| **Model Statistics** |  |  |  |  |
| Map CC (masked) | 0.81 | 0.78 | 0.75 | 0.75 |
| Clash score | 8.36 | 8.26 | 6.72 | 6.04 |
| Molprobity score | 1.99 | 1.86 | 1.80 | 1.71 |
| Cβ deviations | 0 | 0 | 0 | 0 |
| Rotamer outliers (%) | 2.39 | 1.67 | 0 | 0.10 |
| **Ramachandran** |  |  |  |  |
| Outliers | 0 | 0 | 0 | 0 |
| Allowed | 3.72 | 3.62 | 6.67 | 5.59 |
| Favored | 96.28 | 96.38 | 93.33 | 94.41 |
| **Bonds (RMSD)** |  |  |  |  |
| Length (Å) | 0.003 | 0.004 | 0.002 | 0.002 |
| Bond angles | 0.469 | 0.541 | 0.528 | 0.479 |

## Notes

### Competing Interest Statement

The authors have declared no competing interest.

